# Double and triple thermodynamic mutant cycles reveal the basis for specific MsbA-lipid interactions

**DOI:** 10.1101/2023.07.03.547565

**Authors:** Jixing Lyu, Tianqi Zhang, Michael T. Marty, David Clemmer, David H. Russell, Arthur Laganowsky

## Abstract

Structural and functional studies of the ATP-binding cassette transporter MsbA have revealed two distinct lipopolysaccharide (LPS) binding sites: one located in the central cavity and the other at a membrane-facing, exterior site. Although these binding sites are known to be important for MsbA function, the thermodynamic basis for these specific MsbA-LPS interactions is not well understood. Here, we use native mass spectrometry to determine the thermodynamics of MsbA interacting with the LPS-precursor 3-deoxy-D-*manno*-oct-2-ulosonic acid (Kdo)_2_-lipid A (KDL). The binding of KDL is solely driven by entropy, despite the transporter adopting an inward-facing conformation or trapped in an outward-facing conformation with adenosine 5’-diphosphate and vanadate. An extension of the mutant cycle approach is employed to probe basic residues that interact with KDL. We find the molecular recognition of KDL is driven by a positive coupling entropy (as large as -100 kJ/mol at 298K) that outweighs unfavorable coupling enthalpy. These findings indicate that alterations in solvent reorganization and conformational entropy can contribute significantly to the free energy of protein-lipid association. The results presented herein showcase the advantage of native MS to obtain thermodynamic insight into protein-lipid interactions that would otherwise be intractable using traditional approaches, and this enabling technology will be instrumental in the life sciences and drug discovery.

## Introduction

Most Gram-negative bacteria contain outer membrane lipopolysaccharide (LPS) that is crucial for maintaining structural integrity and protection from toxins and antibiotics^1–3^. The **A**TP-**B**inding **C**assette (ABC) transporter MsbA flips an LPS-precursor, lipooligosaccharide (LOS), from the cytosolic leaflet to the periplasmic leaflet of inner membrane, a process powered by the hydrolysis of adenosine triphosphate (ATP). MsbA functions as a homodimer and each subunit consists of a soluble nucleotide-binding domain (NBD) and a transmembrane domain containing six transmembrane helices^4^. The proposed mechanism of MsbA-mediated LOS transportation involves the binding of LOS to the interior binding site and a conformational change from an inward-facing conformation (IF) to an outward-facing conformation (OF).

Like other ABC transporters, the ATPase activity of MsbA can be stimulated in the presence of different substrates, particularly hexaacylated lipid A species.^5–7^ Recent studies have illuminated the location and importance of several LOS binding sites on MsbA^8–11^. The interior binding site is located in the inner cavity, and mutations (R78A, R148A and K299A) engineered to disrupt binding at this site abolish lipid-stimulated ATPase activity and adversely affect cell growth.^8,12^ More recently, the LPS-precursor 3-deoxy-D-manno-oct-2-ulosonic (Kdo)2-lipid A (KDL) was found to bind to an elusive exterior site on MsbA trapped in an OF conformation with adenosine 5’-diphosphate and vanadate^5–7^. Similarly, introducing mutations to disrupt binding at the exterior site also abolishes lipid-induced stimulation of ATPase activity.^10^

In 1984, Fersht and colleagues introduced the biochemistry community to the application of double mutant cycles as means to quantify the strength of intramolecular and intermolecular interactions.^13^ The method has proven to be highly effective in examining pairwise interactions, as demonstrated by its notable application in determining the spatial orientation of potassium channel residues in relation to high-affinity toxin binding.^14^ More generally, the technique has been used to measure the strength and coupling for residues in protein-protein complexes, protein-ligand complexes, and stability of secondary structure.^13–21^ In general, mutant cycles analysis involves measuring the changes in Gibbs free energy for the wild-type protein (P), two single point mutations (PX and PY) and the double mutant protein (PXY) for a given process, such as for protein-protein interactions (for review see ^15^). If residue X and Y are independent of each other, then the Gibbs free energy associated with the double mutant protein will be equal to the sum of changes in Gibbs free energy due to the single mutations relative to the wild-type protein. However, if the Gibbs free energy associated with the structural and functional properties of the double mutant protein differs from the sum of single mutant proteins, then the two residues are energetically coupled or co-operative. The coupling free energy (ΔΔG_int_) is the energy difference between double mutant and two single mutant proteins (see methods). The ΔΔG_int_ values for pairwise interactions in proteins has revealed the contributions of salt bridges (4-20 kJ/mol), aromatic-aromatic interactions (4 kJ/mol), and charge-aromatic interactions (4 kJ/mol) to protein stability.^15,22,23^ Prior work on mutant cycles often employed traditional approaches, but such approaches overlook contributions from conformational changes of the reactants as well as potential changes in the hydration of the complex, including the reacting ligand and the solvent.^16^

To demonstrate the utility of the mutant cycle approach, we highlight two well-known examples. First, the high-affinity interaction between barnase, (an extracellular RNase of *Bacillus amyloliquefaciens*) and barstar (inhibitor of barnase) has been extensively studied by double mutant cycles.^20^ For example, pairwise interactions between residues that are less than seven angstrom in distance (based on crystal structures) have been shown to be co-operative. These interactions were shown to be important for stability of the barnase-barstar complex with coupling energies reaching as high as 7 kcal/mol. Another classical example involves the application of mutant cycles to guide docking and spatial arrangement of a high-affinity peptide inhibitor (scorpion toxin) binding to the Shaker potassium channel.^14^ Of the pairwise interactions that underwent mutant cycle analysis, one pair (R24 from toxin and D431 from channel) in particular displayed an extraordinary coupling energy of 17 kJ/mol. This result indicates the two residues interact in the complex. Despite the absence of a structure of toxin-potassium channel complex, results from the mutant cycle analysis provided a strong constraint positioning the toxin relative to the potassium channel pore-forming region. In summary, these studies demonstrate how mutant cycle analysis can be used to determine the energetics of pairwise interactions, which is important for understanding how these molecular interactions contribute to the overall stability of proteins in complex with other molecules, such as ligands and other proteins.

Native mass spectrometry (MS) is well suited to characterize the interactions between protein and other molecules, especially for membrane proteins.^24–26^ The technique is capable of maintaining non-covalent interactions and native-like structure in the gas phase,^27,28^ essential for studying biochemical interactions with small molecules, such as the binding of drugs, lipids, and nucleotides.^28–35^ In combination with a variable temperature nano electrospray ionization device, native MS has determined the thermodynamics for protein-protein and protein-ligand interactions.^36–41^ For example, the molecular interaction between the signaling lipid 4,5-bisphosphate phosphatidylinositol and Kir3.2 is dominated by a large, favorable change in entropy.^40^ Recently, native MS has been combined with mutant cycles analysis to determine the energetic contribution of pairwise inter-protein interactions for a soluble protein complex.^42,43^ Notably, the coupling energies determined by native MS and isothermal calorimetry are in agreement.^42^ Mutant cycle analysis is also being used to study cardiolipin binding to sites on AqpZ with native MS.^44^

Traditional mutant cycles focus on pairwise interactions, such as two interacting residues in a protein complex.^45^ Single-and double-point mutations along with characterizing their impact on protein stability/assembly enable assessment of the energetic contribution for the pairwise interaction. If the two residues are independent (non-co-operative), then the change in free energy will be equal to the sum of the two single mutations. In contrast, if the two residues are dependent on each other, then the coupling energy is a measure of their co-operativity. Although mutant cycles are often applied to protein-protein interactions, here we extend mutant cycle principles to study membrane protein-lipid interactions. It is established that MsbA has two high-affinity binding sites for the LPS-precursor KDL. Here, we examine each site independently followed by simultaneously probing both KDL sites. At present, there is limited availability of synthetic KDL derivatives, limiting this study to focus on residues that interact with KDL, such as basic residues coordinating the conserved phosphoglucosamine (P-GlcN) of KDL. Despite the limitation of commercially available KDL derivatives, the studies below demonstrate how residues energetically contribute to specific binding, providing insight into the driving forces underlying essential membrane protein-lipid interactions. Recently, we reported results using native MS that reveal conformation-dependent lipid binding affinities to MsbA.^10^ As these measurements were performed at a single temperature, we set out to perform a more detailed thermodynamic analysis to better understand the molecular driving forces that underpin specific MsbA-lipid interactions. Here, we report binding thermodynamics (ΔH, ΔS, and ΔG) for KDL binding to MsbA in IF and OF conformations. These results reveal the unique thermodynamic contributions of MsbA residues that engage KDL. We also report coupling energetics (ΔΔG_int_) for pairwise interactions, including, for the first time, the contributions from coupling enthalpy (ΔΔH_int_) and coupling entropy (Δ(-TΔS_int_)), providing rich molecular insight into specific protein-lipid interactions.

## Results

MsbA residues selected for mutant cycles analyses. MsbA is known to bind KDL either in the inner cavity or at the two exterior sites (**Fig. 1**). For both sites, a series of conserved arginine and lysine residues form specific interactions with the headgroup of KDL. To perform mutant cycles analysis, we introduced single mutations into MsbA to target KDL binding to the interior (MsbA^R78A^ and MsbA^R299A^) and exterior (MsbA^R188A^, MsbA^R238A^, and MsbA^K243A^) sites. More specifically, R78 coordinates one of the characteristic phosphoglucosamine (P-GlcN) substituents of KDL whereas K299 interacts with a carboxylic acid group in the headgroup of KDL. The two P-GlcN constituents of LOS are coordinated by R238 and R188 + K243, respectively. R188 also forms an additional hydrogen bond with the headgroup of KDL. In addition, we prepared double and triple mutants of MsbA for the various residues that were selected for mutagenesis.

**Figure 1.**
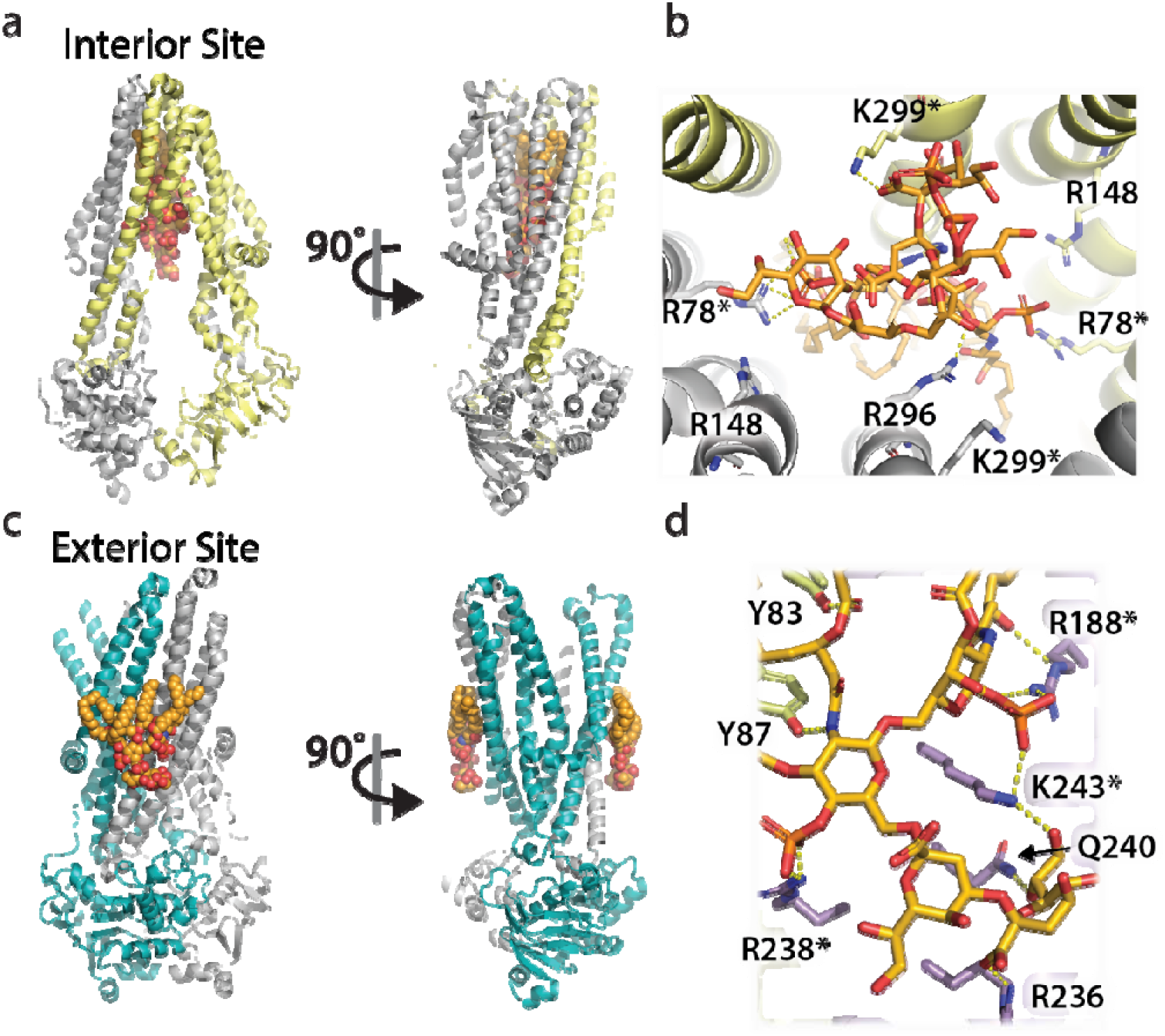
The two distinct LPS binding sites of MsbA and their molecular interactions. a) Two views of LPS bound to the interior site or central cavity of MsbA. The protein shown is also bound to the inhibitor G907 (PDB 6BPL).^9^ The protein and lipid are shown in cartoon and stick representation, respectively. b) Molecular details of the residues interacting with LPS at the interior site. Bonds are shown as dashed yellow lines along with residue labels. c) Two views of the KDL molecules bound to the two exterior binding sites of MsbA that are symmetrically related (PDB 8DMM).^10^ Shown as described in panel A. d) Molecular view of KDL bound to MsbA and shown as described in panel B. The asterisk denotes residues selected for mutant cycle analysis.

### Thermodynamics of MsbA-KDL interactions

We performed titrations to determine the equilibrium binding affinity for MsbA-KDL interactions at four different temperatures (288, 293, 298 and 303 K) (**Fig. 2**). These studies used optimized samples of MsbA that do not contain any co-purified LPS (for details see ^10^). The transporter was stable at the selected temperatures. For example, binding of KDL to MsbA was enhanced at higher temperatures (**Fig. 2a**), indicating a favorable entropy for the interaction. For a given temperature, the mass spectra from the titration series were deconvoluted and equilibrium dissociation constants (K_D_) were determined for MsbA binding up to three KDL molecules (**Fig. 2b, Supplementary** Fig. 1 **and Supplementary Table 1**). It is important to note that, unlike traditional approaches that struggle to distinguish between free protein from that bound to ligand,^46^ native MS can resolve different ligand bound states, including the free concentration of protein and free concentration of ligand(s), in a single mass spectrum.^36,47–49^ Notably, the native MS approach has been cross validated using isothermal calorimetry and surface plasmon resonance.^36,47,48^ Interestingly, van’ t Hoff analysis showed a non-linear trend for three KDL binding reactions (**Fig. 2c**), indicating that over the selected temperature range, heat capacity is not constant.^50^ The nonlinear form of the van’t Hoff equation enabled us to determine the ΔH and change in heat capacity (ΔC_p_) at a reference temperature of 298CK (**Fig 2c-d**). In this case, ΔG was calculated directly from K_D_ values, and entropy (ΔS) was back calculated using both ΔH and ΔG. ΔG values for binding KDL_1-2_ range from -32.0 ± 0.1 to -35.2 ± 0.1 kJ/mol. The binding reaction has a positive ΔC_p_ that alters the thermodynamic parameters at different temperatures. At the lowest temperature, KDL binding is driven by favorable enthalpy (-36 ± 12 to -43 ± 7 kJ/mol) with a small entropically penalty (-TΔS, 2 ± 11 to 12 ± 7 kJ/mol at 288 K). In contrast, KDL binding at higher temperatures displays a large, favorable entropy (-TΔS, -123 ± 12 to -146 ± 7 kJ/mol at 303 K) that compensates a large enthalpic barrier (86 ± 12 to 112 ± 7 kJ/mol). These results highlight the role of entropy in KDL binding to MsbA that may stem from solvent reorganization.

**Figure 2.**
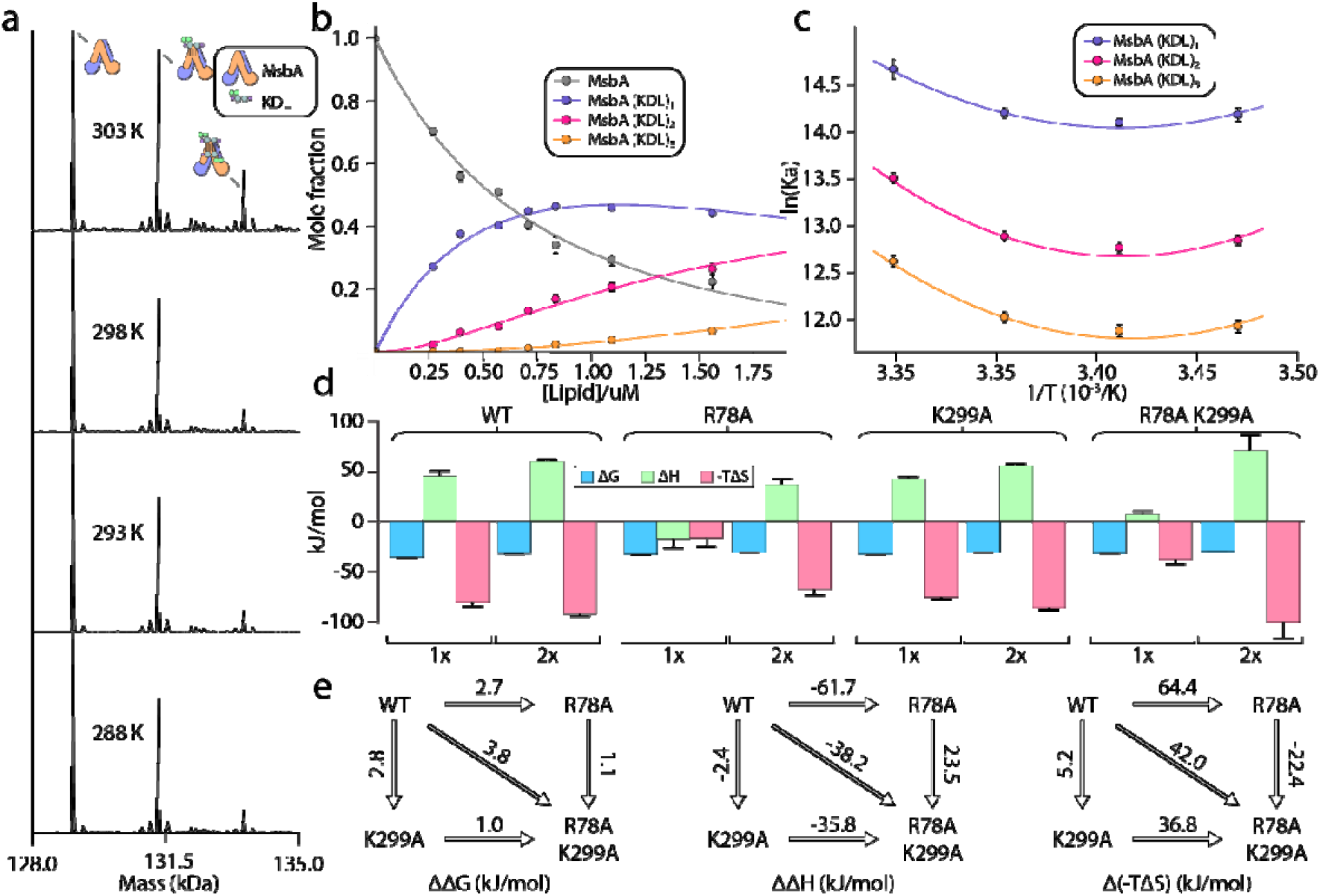
Thermodynamics of KDL binding at the interior site to wild-type and mutant MsbA. a) Representative deconvoluted native mass spectra of 0.39 μM wild-type MsbA in C10E5 and in the presence of 0.6 μM KDL recorded at different solution temperatures. b) Plot of mole fraction of MsbA (KDL)_0-3_ determined from titration of KDL (dots) at 298 K and resulting fit from a sequential ligand binding model (solid line, R^2^ = 0.99). c) van’ t Hoff plot for MsbA(KDL)_1-3_ and resulting fit of a nonlinear van’ t Hoff equation. d) Thermodynamics for MsbA and mutants (MsbA^R78A^, MsbA^K299A^ and MsbA^R78A,K299A^) binding KDL at 298 K. e) Mutant cycles for MsbA and mutants with (from left to right) ΔΔG (mutant minus wild-type), ΔΔH and Δ(-TΔS) values indicated over the respective arrows. Shown are values at 298K. Reported are the average and standard deviation from repeated measurements (n1=13).

### KDL binding to the interior binding site of MsbA

We next determined the thermodynamics of KDL binding to MsbA containing single and double mutations at the interior binding site (**Fig 2d**). R78 of each subunit interacts with one of the P-GlcN moieties of LPS whereas one of the K299 residues interacts with a carboxylic acid group of LPS molecule in the inner cavity. Therefore, introducing the R78A mutation will impact symmetrically equivalent binding sites. MsbA^R78A^ showed a reduction in binding KDL_1-2_ with ΔG ranging from -30.2 ± 0.2 to -32.5 ± 0.2 kJ/mol. At 298K, KDL binding is enthalpically and entropically favorable whereas binding of the second KDL is similar to the wild-type protein (**Fig. 2d**). The binding thermodynamics for MsbA^K299A^ is reminiscent of the wild-type protein with a large, favorable change in entropy (-TΔS, -75 ± 2 to -86 ± 1 kJ/mol at 298 K) and unfavorable enthalpy (43 ± 2 to 53 ± 1 kJ/mol) (**Fig. 2d**). The double mutant MsbA^R78A,K299A^ protein shows a reduction in opposing entropic and enthalpic terms leading to an increase in ΔG by ∼4 kJ/mol relative to wild-type MsbA (**Fig. 2d**). Mutant cycle analysis indicates a coupling energy (ΔΔG_int_) of 1.7 *±* 0.4 kJ/mol that contributes to the stability of KDL-MsbA complex (**Fig. 2e** and **Supplementary Table 3**). More generally, ΔΔ with a positive sign means favorable cooperation. Interestingly, the coupling enthalpy (ΔΔH_int_ of -26 ± 15 kJ/mol) and coupling entropy (Δ(-TΔS)_int_ of 28 ± 15 kJ/mol at 298K) indicating that these residues contribute to KDL binding through an entropy driven process that overcomes an enthalpic barrier (**Fig. 2e** and **Supplementary Table 3**).

### KDL binding to the exterior binding site of MsbA

The recently discovered exterior KDL binding site^10^ located on the cytosolic leaflet of inner membrane has not been thoroughly investigated, prompting us to characterize this site by a triple mutant cycle (**Fig 3**). We first investigated R188 and K243, residues that both interact with one of the P-GlcN moieties of LOS. Like mutants targeting the interior LPS binding site, introducing mutants at the exterior site will impact binding at the two exterior sites. Both MsbA^R188A^ and MsbA^K243A^ single mutants marginally weakened the interaction by about 2 kJ/mol (**Fig 3a**). Enthalpy and entropy for KDL binding MsbA^R188A^ and MsbA^K243A^ was largely similar to the wild-type protein (**Fig 3a**). However, the R238A mutation significantly weakened the interaction with KDL, increasing ΔG by nearly 5 kJ/mol compared to the wild-type transporter (**Fig 3b** and **Supplementary Table 3**) and resulted in an distinct thermodynamic pattern with negative enthalpy changes for both the first and second KDL binding events (**Fig 3a**). ΔG for MsbA^R188A,K243A^ was comparable to the K243A single mutant form of the protein (**Fig 3a**). The positive coupling energy of 3.2 ± 0.4 kJ/mol with contributions from a coupling enthalpy of 19 ± 11 kJ/mol and a coupling entropy of -16 ± 12 kJ/mol at 298K (**Fig 3b** and **Supplementary Table 3**). Combining mutation R238A with R188A, MsbA^R188A,R238A^ decreased ΔH by 57 kJ/mol at the cost of increasing -TΔS by 64 kJ/mol at 298K (**Fig 3b** and **Supplementary Table 3**). The coupling energy for R188A and R238A is approximately zero as a result of equal coupling enthalpy and entropy of different signs. Compared to the wild-type protein, MsbA^R238A,K243A^ results in an inversion of the thermodynamic signature with binding now being driven by enthalpy. More specifically, this inversion is accompanied by ΔΔH and Δ(-TΔS) of -93 ± 7 kJ/mol and 99 ± 7 kJ/mol at 298K (**Fig 3b** and **Supplementary Table 3**). Again, the coupling enthalpy and entropy (at 298K) of equal magnitude but opposite signs give rise to a coupling energy of zero for R238A and R243A (**Fig 3b** and **Supplementary Table 3**). Introduction of the R188A mutation into MsbA^R238A,K243A^, results in reversal of the thermodynamic signature to mirror that of MsbA^R188A,K243A^ (**Fig 3a**) . The coupling energy, coupling enthalpy, and coupling entropy for R188A, R238A and R243A are 3.4 ± 0.5 kJ/mol, 100 ± 16 kJ/mol, and -97 ± 16 kJ/mol at 298K (**Supplementary Table 4**), respectively. Taken together, these results demonstrate KDL binding to MsbA is sensitive to mutations at both the interior and exterior sites.

**Figure 3.**
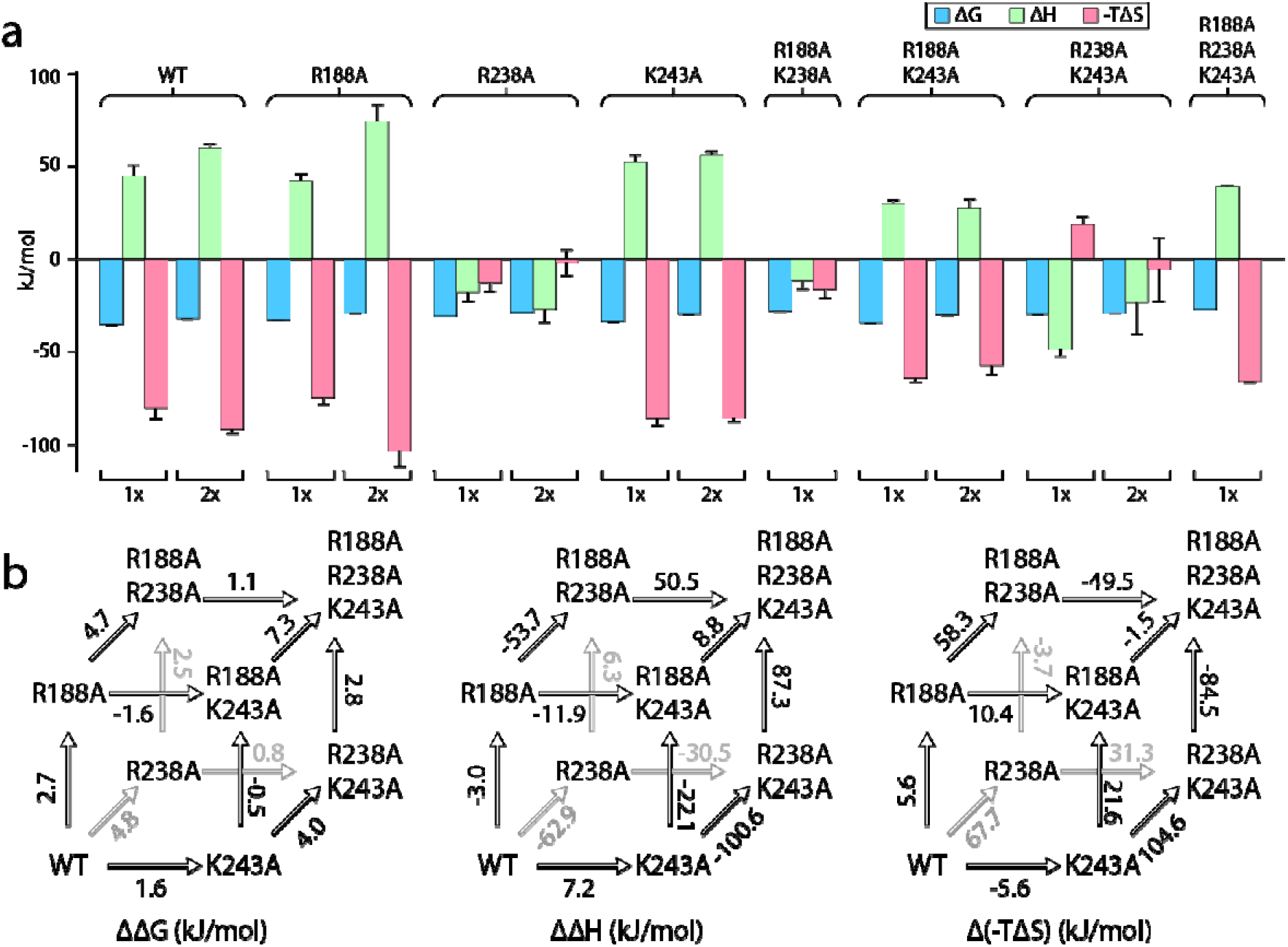
Triple mutant cycle analysis of the exterior LPS binding site of MsbA. a) Thermodynamics for MsbA and mutants (MsbA^R188A^, MsbA^R238A^, MsbA^K243A^, MsbA^R188A,R243A^, MsbA^R188A,K243A^, MsbA^R238A,R243A^, and MsbA^R188A,R238A,K299A^) binding KDL at 298 K. b) Triple mutant cycles for MsbA and mutants with (from left to right) ΔΔG, ΔΔH and Δ(-TΔS) values indicated over the respective arrows. Shown are values at 298K. Reported are the average and standard deviation from repeated measurements (n1=13).

### Dissecting KDL binding to the interior and exterior site(s) of MsbA

An open question is if the interior and exterior LOS binding sites of MsbA are allosterically coupled? We focused on the R188A and K299A mutants located at the exterior and interior binding sites, respectively. Results for both single mutants were presented above. MsbA containing the R188A and K299A mutations drastically reduced the binding of KDL (**Fig 4a**). The ΔG for MsbA^R188A,K299A^ increased by more than 6 kJ/mol compared to the wild-type protein (**Fig 4b**). This approximately doubles compared to MsbA containing either of the single point mutations. Mutant cycle analysis revealed a negative coupling energy of -1.1 ± 0.4 kJ/mol that partitioned into a coupling enthalpy of -36 ± 13 kJ/mol and coupling entropy of 34 ± 13 kJ/mol at 298K (**Fig. 4b** and **Supplementary Table 3**). In short, mutations at either LOS binding site have a negative impact on binding that is accompanied by a gain in both favorable entropy and unfavorable enthalpy.

**Figure 4.**
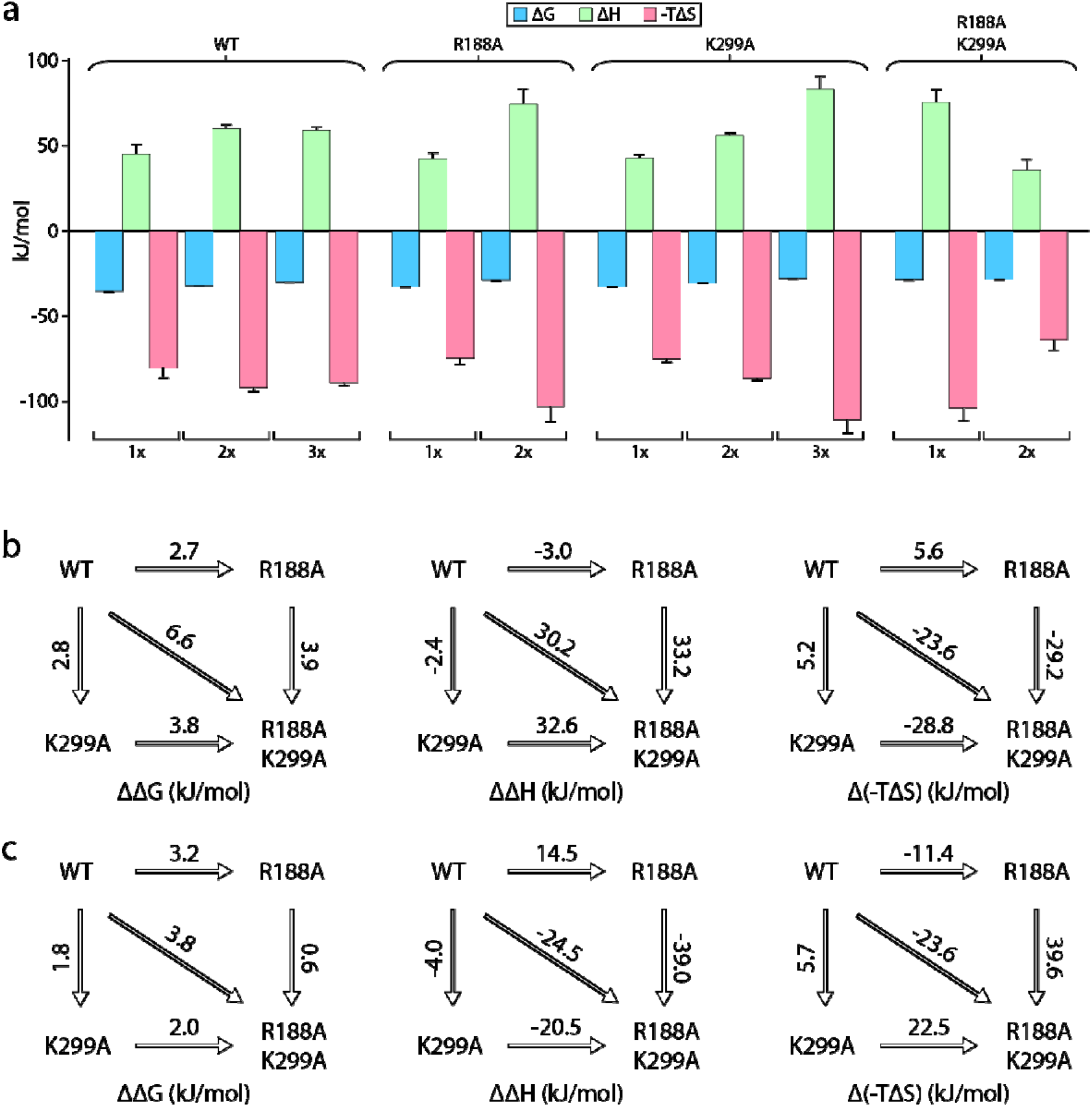
Mutant cycle of MsbA residues located within the interior and exterior LOS bind sites. . a) Thermodynamic signatures for MsbA and mutants binding KDL at 298 K. b-c) Double mutant cycle analysis for R188 and K299. Shown are results for the first (panel b) and second (panel c) KDL binding to MsbA. Shown from left to right is ΔΔG, ΔΔH and Δ(-TΔS) and the values indicated over the respective arrows at 298K. Reported are the average and standard deviation from repeated measurements (n1=13).

### Mutant cycle analysis of KDL binding to vanadate-trapped MsbA

As MsbA, like other ABC transporters, is highly dynamic, we sought to trap the transporter in an OF conformation using ADP and vanadate to interrogate binding at the exterior lipid binding site. We characterized the binding of KDL to vanadate-trapped MsbA and proteins containing single R188A, R238A, and K243A mutations. Here, we focused on the binding of the first and second lipid, since MsbA has two, symmetrically related KDL binding sites in the open, OF conformation. Thermodynamics of MsbA(KDL)_1-2_ binding is like the non-trapped transporter, wherein entropy (-TΔS ranging from -58 ± 1 to -69 ± 1 kJ/mol at 298K) is more favorable than a positive enthalpic term (ΔH ranging from 22 ± 1 to 35 ± 1 kJ/mol) (**Fig. 5a**). The single mutant proteins (MsbA^R188A^, MsbA^R238A^, and MsbA^K243A^) showed a slight increase in ΔG (at most 5 kJ/mol) (**Fig. 5a**). Notably, we found MsbA^R238A^ and MsbA^K243A^ had about a four-fold increase in ΔH and favorable entropy was about two-fold higher (**Fig. 5a**). Double mutant cycle analysis of the pairwise mutants revealed a positive coupling energy of ∼2 kJ/mol for MsbA binding one and two KDLs (**Fig. 5b** and **Supplementary Table 7**). Focusing on the first KDL binding event, the coupling enthalpy and coupling entropy at 298K for R188 and K238 was 89 ± 7 kJ/mol and -87 ± 7 kJ/mol, respectively (**Fig. 5b** and **Supplementary Table 7**). Likewise, R238 and K243 showed 129 ± 11 kJ/mol of coupling enthalpy and -127 ± 11 kJ/mol of coupling entropy at 298K (**Fig. 5b** and **Supplementary Table 7**). However, the R188 and K243 pair revealed a relatively low coupling enthalpy and coupling entropy at 298K of 3.5 ± 7 kJ/mol and 2 ± 7 kJ/mol, respectively (**Fig. 5b** and **Supplementary Table 7**). These results highlight the importance of entropic and enthalpic contributions that underpin specific lipid binding sites.

**Figure 5.**
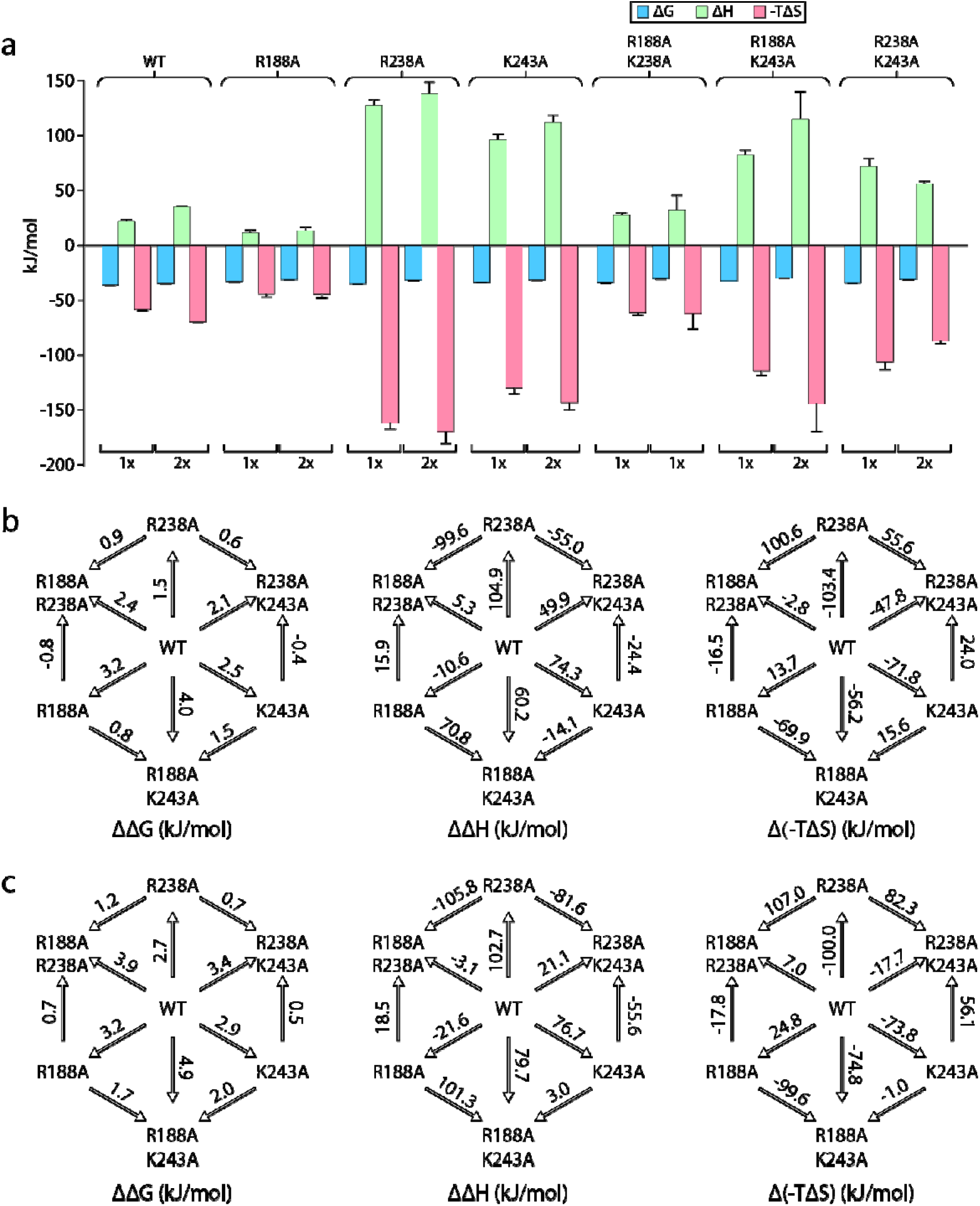
Double mutant cycles reveal thermodynamic insight into KDL binding vanadate-trapped MsbA. a) Thermodynamic signatures for MsbA and mutants binding KDL at 298 K. b-c) Double mutant cycle analysis for pairs of R188, R238, and K243 with a total of three combinations. Shown are results for the first (panel b) and second (panel c) KDL binding to MsbA trapped in an open, OF conformation with ADP and vanadate. Within each panel, ΔΔG, ΔΔH and Δ(-TΔS) are shown from left to right and their values at 298K indicated over the respective arrows. Reported are the average and standard deviation from repeated measurements (n1=13).

## Discussion

Thermodynamics provide unique insight into the molecular forces that drive specific MsbA-KDL interactions. A recurring thermodynamic strategy for specific KDL-MsbA interactions is a large, favorable entropic term that opposes a positive enthalpic value. The human G-protein-gated inward rectifier potassium channel (Kir3.2) also used a similar thermodynamic strategy to engage phosphoinositides (PIPs).^40^ The large, positive entropy could stem from solvent reorganization of the lipid with a carbohydrate containing headgroup, and desolvation of hydrated binding pockets on the membrane protein. The release of ordered solvent to the bulk solvent would contribute favorably to entropy. These experiments are performed in detergent and reorganization of detergent may also play a role. Previous work has shown soluble protein-ligand interactions can be driven by a large, positive entropy term that outweighs a large, positive enthalpic penalty.^51–54^ In these cases, the reaction is mainly driven by conformational entropy originating in enhanced protein motions. However, it is unclear if the conformational dynamics of MsbA are enhanced when bound to KDL.

Most of the van’ t Hoff plots followed non-linear trends, indicating C_p_ is not constant over the selected temperature range.^50^ In nearly all cases, a positive ΔC_p_ was observed that ranged in value from 4 to 12 kJ/mol·K (**Supplementary Table 2, 6**). Solvation of polar groups in aqueous solvent has been ascribed to positive heat capacities whereas the collapse of apolar residues from their solvated state is accompanied by a negative change in heat capacity.^50,55^ Reorganization of the hydrated, polar headgroups of KD is consistent with the positive heat capacity observed here. However, change in heat capacity could also be ascribed to temperature-dependent conformational changes in MsbA and/or KDL. Notably, vanadate-trapped MsbA locked in an open, OF conformation should be less conformationally dynamic than the apo protein, which is known to adopt a number of open, IF conformations where the NBDs are separated by different distances. Similar positive heat capacities were observed for the different conformations, suggesting the dynamics of MsbA marginally contribute to the observed non-linear trends. Notably, the headgroup of KDL is nestled in a hydrophilic, basic patch of MsbA in the open, OF conformation. Similarly, the headgroup of PIP binds a hydrophilic, basic pocket in Kir3.2. These hydrophilic patches will be highly solvated, which will be desolvated upon binding lipids contributing favorably to entropy.

Thermodynamics of MsbA-lipid interactions contrast those observed for a different membrane protein. Phospholipid binding to the bacterial ammonia channel (AmtB) were largely driven by enthalpy and, in most cases, entropy was unfavorable.^56^ Another interesting observation for AmtB-lipid interactions was significant enthalpy-entropy compensation for each sequential lipid binding event. Here, enthalpy-entropy compensation is not as pronounced. This result may reflect the much higher-affinity and specific MsbA-KDL interactions compared to the weaker AmtB-lipid interactions, sometimes referred to as non-annular lipids.^24^ Moreover, we have focused the titration here to characterize the binding of the first three KDL molecules to MsbA. While we can’t rule out that the resolved lipid bound states of MsbA represent binding of lipid to one or multiple site(s) on the transporter, the mole fraction plots are suggestive of binding to distinct sites, i.e., smooth inflections. In a previous study,^40^ we observed abnormal binding curves for some PIPs binding to Kir3.2 that we rationalized by the presence of high-affinity binding and low-affinity binding sites. A revised equilibrium binding model including the two-site model dramatically improved the fits, leading to dissection of at least two lipid binding sites. Further studies are warranted to better understand the binding sites of KDL to MsbA in different conformations.

Results of this study begin to draw a connection between LPS binding at the interior and exterior sites of MsbA. It is presently thought that flipping of LOS occurs at interior MsbA site, and the exterior LOS binding site enables feedforward activation, wherein binding of LOS and precursors thereof stimulates ATPase activity.^8–10,57–60^ It is also thought that binding of LOS and ATP promotes dimerization of the NBDs. Here, we find mutations at either the interior or exterior sites have a direct impact of KDL binding to MsbA, which under these conditions is presumably adopting an open, IF conformation. Of the mutant proteins, MsbA containing single mutations (MsbA^R188A,K299A^) at both LOS binding sites resulted in the greatest change in ΔG. This result implies that these sites are allosterically coupled and further investigation is warranted to better understand how the exterior LOS binding sites influence MsbA dynamics.

A defining feature of this work is the use of mutant cycles to not only characterize specific membrane protein-lipid interactions but define the coupling energies of specific residue-lipid interactions in terms of enthalpic and entropic contributions. Traditionally, mutant cycles have been used to understand pairwise interactions of residues, such as in protein-protein complexes, in terms of coupling free energy. Here, we extend mutant cycles to understand how pairs of residues contribute to specific MsbA-KDL interactions. Double mutants targeting the interior site reveal a positive coupling energy of nearly 2 kJ/mol for R78 and K299. These stabilize the MsbA-KDL complex largely through nearly 17 kJ/mol of favorable coupling entropy, which outweighs a negative coupling enthalpy. This phenomenon extends to nearly all mutant cycles investigated in this work, even when the transporter is trapped with vanadate. The largest coupling energy is observed from the triple mutant cycle of R188A, R238A and R243A, which again stabilization of the complex was achieved via favorable coupling entropy. While we focused on results at 298K, the coupling energetics among these three residues show 3.4 ± 0.5 kJ/mol. Taken together, mutant cycle analysis reveals that entropy drives high-affinity KDL binding to MsbA and solvent reorganization contributes to KDL binding (**Figure 6**). There are many factors that contribute to the change in entropy of the system, beyond solvation entropy, and deciphering the entropic contributions of the various components warrants additional studies.

**Figure 6.**
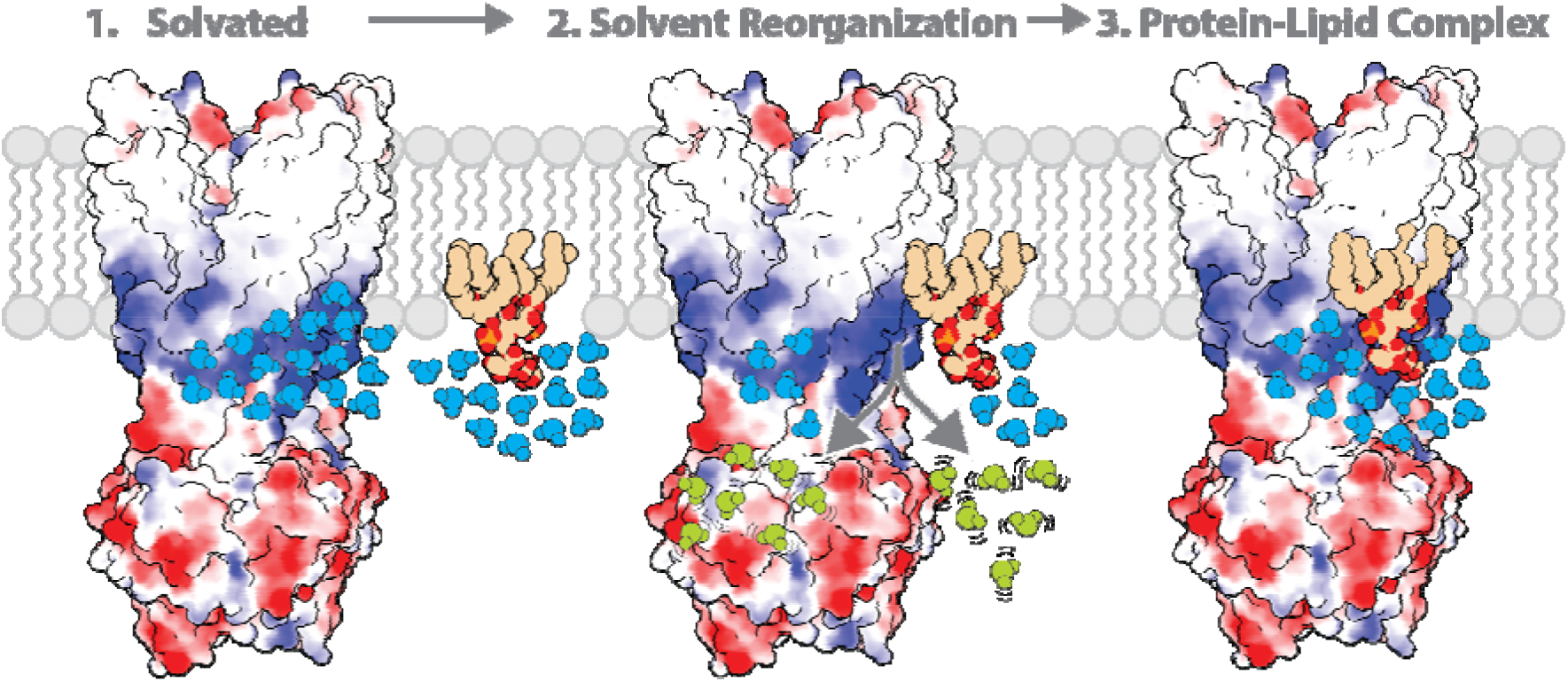
The role of solvent in contributing to the molecular recognition of membrane protein-lipid complexes. The lipid headgroup and binding pocket (basic patch illustrated in blue) on the membrane protein are solvated. The ordered solvent (shown in light blue) is then displaced upon lipid binding the membrane protein leading to solvent reorganization. The displacement of ordered solvent (show in light green) contributes to favorable entropy. This process enables the formation of a high affinity, stable membrane protein-lipid complex.

While the use of mutant cycles was prominent a few decades ago, native mass spectrometry opens new opportunities to revisit the classical approach, diving deeper into the energetics of non-covalent interactions, such as dissecting energetics in terms of enthalpic and entropic contributions. Native MS coupled with a variable-temperature nanoelectrospray ionization (nESI) apparatus^41,47^ has been used to ascertain equilibrium binding constants and thermodynamic properties of protein-protein and protein-ligand interactions. The results obtained align closely with those obtained through other biophysical techniques, such as isothermal titration calorimetry (ITC) and surface plasmon resonance (SPR).^48,56,61^ The approach has also uncovered that specific protein-lipid interactions can allosterically modulate other interactions with protein, lipid, and drug molecules. ^33,34,48,49,62,63^ More recently, native MS has proved useful in dissecting the thermodynamics of individual nucleotide binding events to GroEL, a 801 kDa tetradecameric chaperonin.^39^ In contrast to traditional approaches, such as ITC and SPR, native MS can resolve and dissect individual binding events enabling the measurement of binding thermodynamics, which is of paramount importance to understanding the molecular driving forces of non-covalent interactions.

In summary, we demonstrate the utility of native MS to determine the thermodynamic origins of specific KDL-MsbA interactions. Combined with the classical mutant cycle approach,^13^ the thermodynamic contribution of specific interactions with lipids is illuminated. More specifically, MsbA binding KDL is solely driven by entropy, which overcomes an enthalpic penalty. A similar thermodynamic strategy was also observed for Kir3.2-PIP interactions, where entropy plays a central role in the wild-type channel recognizing PIP.^40^ It is tempting to speculate that favorable entropy is a common theme enabling membrane proteins to specifically engage carbohydrate-containing lipids. We envision thermodynamics and mutant cycles will be invaluable in not only better understanding high-affinity lipid binding sites but also in the development of inhibitors, such as those that may target specific protein-lipid binding site(s). In closing, these studies provide deeper insight into the thermodynamic strategies membrane proteins exploit to achieve high-affinity lipid binding site(s).

## Methods

### MsbA expression constructs

MsbA and mutants were essentially expressed and purified as previously described.^10^ In detail, the MsbA gene (from *Escherichia coli* genomic DNA) was amplified by polymerase chain reaction (PCR) using Q5 High-Fidelity DNA Polymerase (New England Biolabs, NEB) and subcloned into a modified pCDF-1b plasmid (Novagen) resulting in expression of MsbA with an N-terminal TEV protease cleavable His6 fusion protein. Primers for generating mutations for MsbA were designed using the online tool NEBaseChanger (NEB) and carried out using the KLD enzyme mix (NEB) as described by the manufacturer. All plasmids were confirmed by DNA sequencing.

### Protein expression and purification

MsbA expression plasmids were transformed into *E. coli* (DE3) BL21-AI competent cells (Invitrogen). A single colony was picked and used to inoculate 50 mL LB media to be grown overnight at 37 °C with shaking. The overnight culture was used to inoculate to terrific broth (TB) media and incubated at 37 °C until the OD_600nm_ *≈* 0.6-1.0. After which, the cultures were induced with final concentration of 0.5 mM IPTG (isopropyl β-D-1-thiogalactopryanoside) and 0.2% (w/v) arabinose. After overnight expression at 25 °C, the cultures were harvested at 4000 x g for 10 minutes and the resulting pellet was resuspended in lysis buffer (20 mM Tris, 300 mM NaCl and pH at 7.4 at room temperature). The resuspended cells were centrifuged, and the pellet was then resuspended in lysis buffer. Cells were lysed by four passages through a Microfluidics M-110P microfluidizer operating at 25,000 psi with reaction chamber emersed in an ice bath. The lysate was clarified by centrifugation at 20,000 x g for 25 minutes and the supernatant was centrifuged at 100,000 x g for 2 hours to pellet membranes. Resuspension buffer (20 mM Tris, 150 mM NaCl, 20% (v/v) glycerol, pH 7.4) was used to homogenize the resulting pellet and 1% (m/v) DDM was added for protein extraction overnight at 4 °C. The extraction was centrifuged at 20,000 x g for 25 minutes and the resulting supernatant was supplemented with 10 mM imidazole and filtered with a 0.45 µm syringe filter prior to purification by immobilized metal affinity chromatography. The extraction containing solubilized MsbA was loaded onto a column packed with 2.5 mL Ni-NTA resin pre-equilibrated in NHA-DDM buffer (20 mM Tris, 150 mM NaCl, 10 mM imidazole, 10% (v/v) glycerol, pH 7.4 and supplemented with 2x the critical micelle concentration (CMC) of DDM). After the loading, the column was washed with 5 column volumes (CV) of NHA-DDM buffer, 10 CV of NHA-DDM buffer supplemented with additional 2% (w/v) nonyl-ß-glucoside (NG), and 5 CV of NHA-DDM buffer. The immobilized protein was eluted with the addition of 2 CV of NHB-DDM buffer (20 mM Tris, 150 mM NaCl, 250 mM imidazole, 10% (v/v) glycerol, 2 x CMC of DDM, pH 7.4). The eluted MsbA was pooled and desalted using HiPrep 26/10 desalting column (GE Healthcare) pre-equilibrated in desalting buffer (NHA-DDM with imidazole omitted). TEV protease (expressed and purified in-house) was added to the desalted MsbA sample and incubated overnight at room temperature. The sample was passed over a pre-equilibrated Ni-NTA column and the flow-through containing the cleaved MsbA protein was collected. The pooled protein was concentrated using a centrifugal concentrator (Millipore, 100 kDa) prior to injection onto a Superdex 200 Increase 10/300 GL (GE Healthcare) column equilibrated with 20 mM Tris, 150 mM NaCl, 10% (v/v) glycerol and 2x CMC C_10_E_5_. Peak fractions containing dimeric MsbA were pooled, flash frozen in liquid nitrogen, and stored at -80 °C prior to use.

### Preparation of MsbA for native MS studies

MsbA samples were incubated with 20 µM copper (II) acetate, to saturate the N-terminal metal binding site,^10^ prior to buffer exchange using a centrifugal buffere exchange device (Bio-Spin, Bio-Rad) into 200 mM ammonium acetate supplemented with 2 x CMC of C_10_E_5_. To prepare vanadate-trapped MsbA, ATP and MgCl_2_ were added to MsbA at a final concentration of 10mM. After incubation at room temperature for 10 minutes, a freshly boiled vanadate solution (pH 10) was added to reach final concentration of 1 mM followed by incubation at 37 °C for an additional 10 minutes. The sample was then buffer exchanged as described above.

### Native Mass Spectrometry

Samples were loaded into gold-coated glass capillaries made in-house^64^ and introduced into at a Thermo Scientific Exactive Plus Orbitrap with Extended Mass Range (EMR) using native electrospray ionization source modified with a variable temperature apparatus.^41^ For native mass analysis, the instrument was tuned as follow: source DC offset of 10 V, injection flatapole DC to 8.0 V, inter flatapole lens to 4, bent flatapole DC to 3, transfer multipole DC to 3 and C trap entrance lens to 0, trapping gas pressure to 6.0 with the in-source CID to 65.0 eV and CE to 100, spray voltage to 1.70 kV, capillary temperature to 200 °C, maximum inject time to 200 ms. Mass spectra were acquired with a setting of 17,500 resolution, microscans set to 1 and averaging set to 100.

### Determination MsbA-lipid equilibrium binding constants

KDL (Avanti) stock solution was prepared by dissolving lipid powder in water. The concentration of MsbA and KDL were determined by a DC protein assay (BioRad) and phosphorus assay, respectively.^65,66^ MsbA was incubated with varying concentrations of KDL before loading into a glass emitter and mounted on a variable-temperature electrospray ionization (vT-ESI) source.^41^ Samples were incubated in the source for two minutes at the desired temperature before data acquisition. All titration data were collected in triplicate. At a given temperature, the mass spectra were deconvoluted using Unidec^67^ and the peak intensities for apo and KDL-bound species were determined and converted to mole fraction. The sequential ligand binding model was applied to determine the mole fraction of each species in measurement:

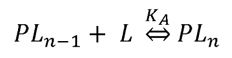

Where:

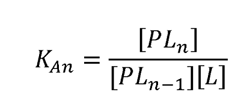

To calculate the mole fraction of a particular species:^56^

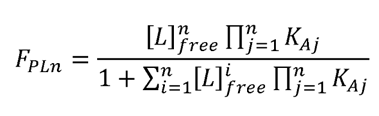

For each titrant in the titration, the free concentration of lipid was computed as follows:

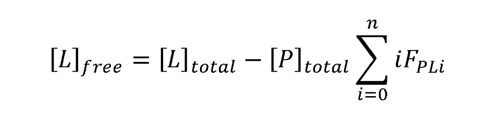

The sequential ligand binding model was globally fit to the mole fraction data by minimization of pseudo-x^2^ function:

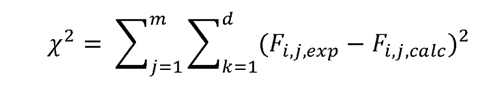

where *n* is the number of bound ligands and *d* is the number of the experimental mole fraction data points.

Van’t Hoff analysis^68^ was applied to determine the Gibbs free energy change (ΔG), enthalpy change (ΔH) and entropy change (ΔS) based on the equation:

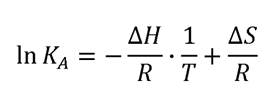

For non-linear trends, the non-linear form of the Van’t Hoff equation was applied to determine the thermodynamic parameters:^69^

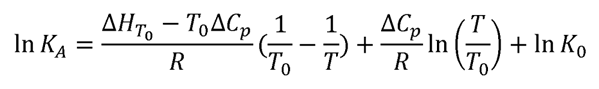

where K_A_ is the equilibrium association constant, K_0_ is the equilibrium association constant at the reference temperature (T_0_), ΔH_TO_ is the standard enthalpy at T_0_, ΔC_p_ is the change in heat capacity at constant pressure, and R is the universal gas constant.

### Mutant cycle analysis

If the two mutated residues are interacting, then the coupling free energy (ΔΔG_int_) will not be 0 and the value may be positive or negative depending upon whether the interactions between mutated residues enhance or weaken the functional property measured.^70^ ΔΔG_int_ can be computed given the change in Gibbs free energy for the wild-type protein (P), two single mutants (PX and PY), and double mutant (PXY) as follows:

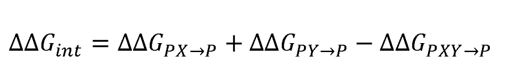

where 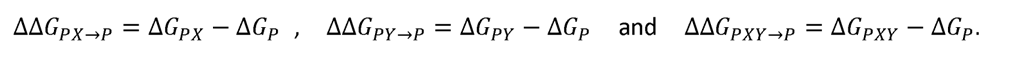 Analogously, the contributions from coupling enthalpy (ΔΔH_int_) and coupling entropy (Δ(-TΔS_int_)) can be computed as follows:

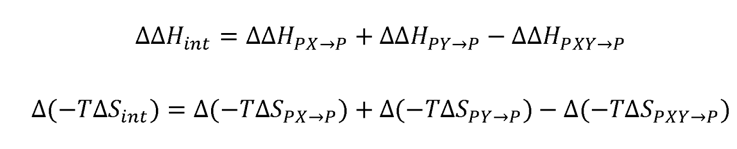

where T is temperature in K. As an example, 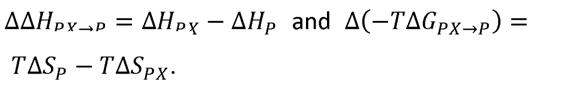

## Acknowledgements

This work was supported by National Institutes of Health (NIH) under grant numbers (DP2GM123486, R01GM121751, R01GM139876, R01GM138863 and RM1GM145416 to A.L.; P41GM128577 to D.R.; and R35GM128624 to M.M.).

## Author Contributions

J.L., M.M., and A.L. designed the research on mutant cycle analysis. J.L. and T.Z. expressed and purified MsbA. J.L. performed mass spectrometry experiments. J.L. and T.Z. carried out the functional assays. J.L., T.Z., M.M., D.C., D.R., and A.L. analyzed the data. J.L. and A.L. wrote the manuscript with input from the other authors.

## Materials & Correspondence

Correspondence and requests for materials should be addressed to AL.

