## Supplementary Information for "Double and triple thermodynamic mutant cycles reveal the basis for specific MsbA-lipid interactions"

**Supplementary Figures**

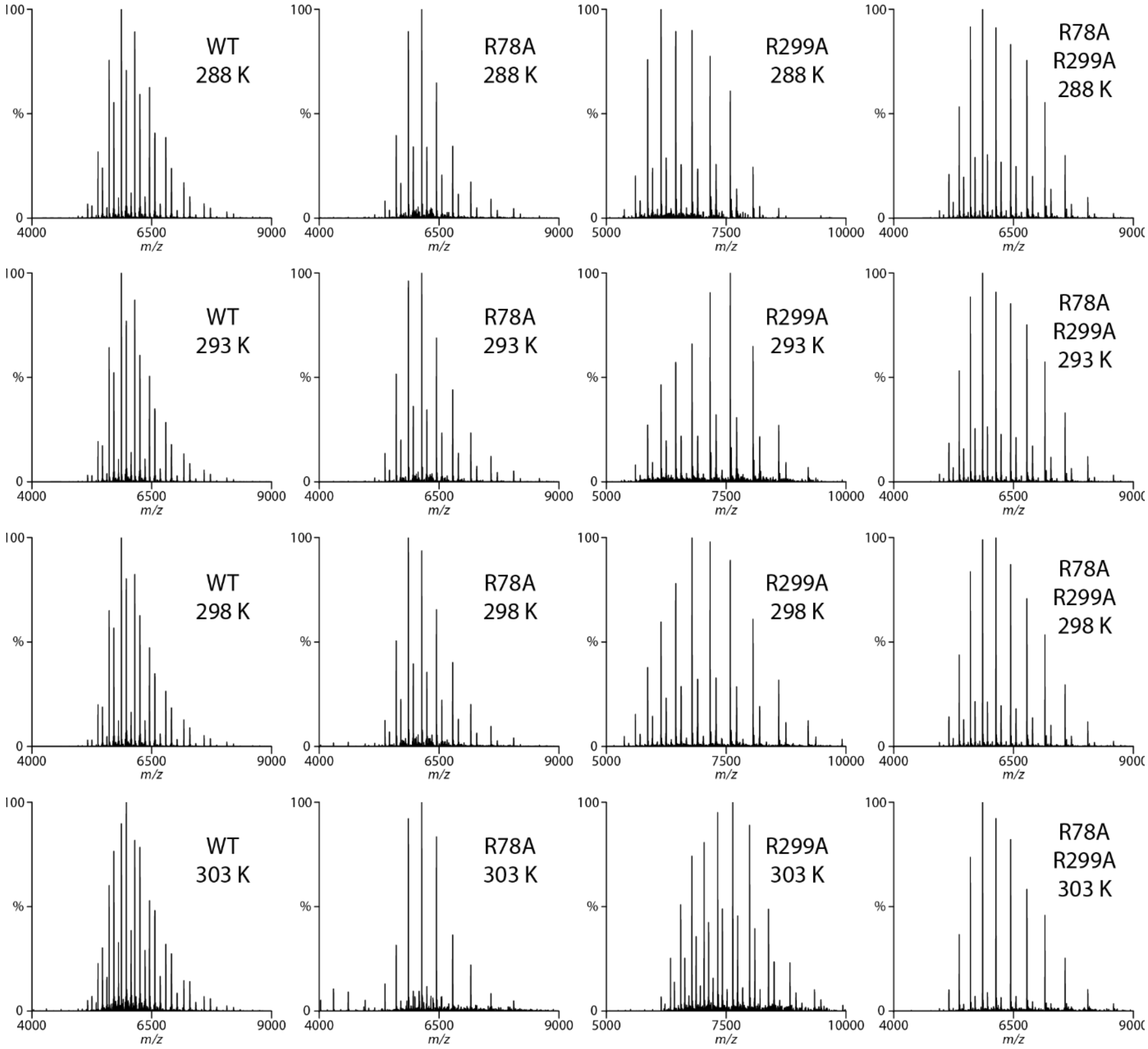


**Supplementary Figure 1**. **Representative native mass spectra of wild-type and mutant MsbA in the presence of 0.8 µM KDL.** The temperature is provided in the inset. The concentration of MsbA, MsbA$^{R78A}$, MsbA$^{R299A}$ and MsbA$^{R78A,K299A}$ is 0.39, 0.36, 0.33, and 0.83 µM, respectively.

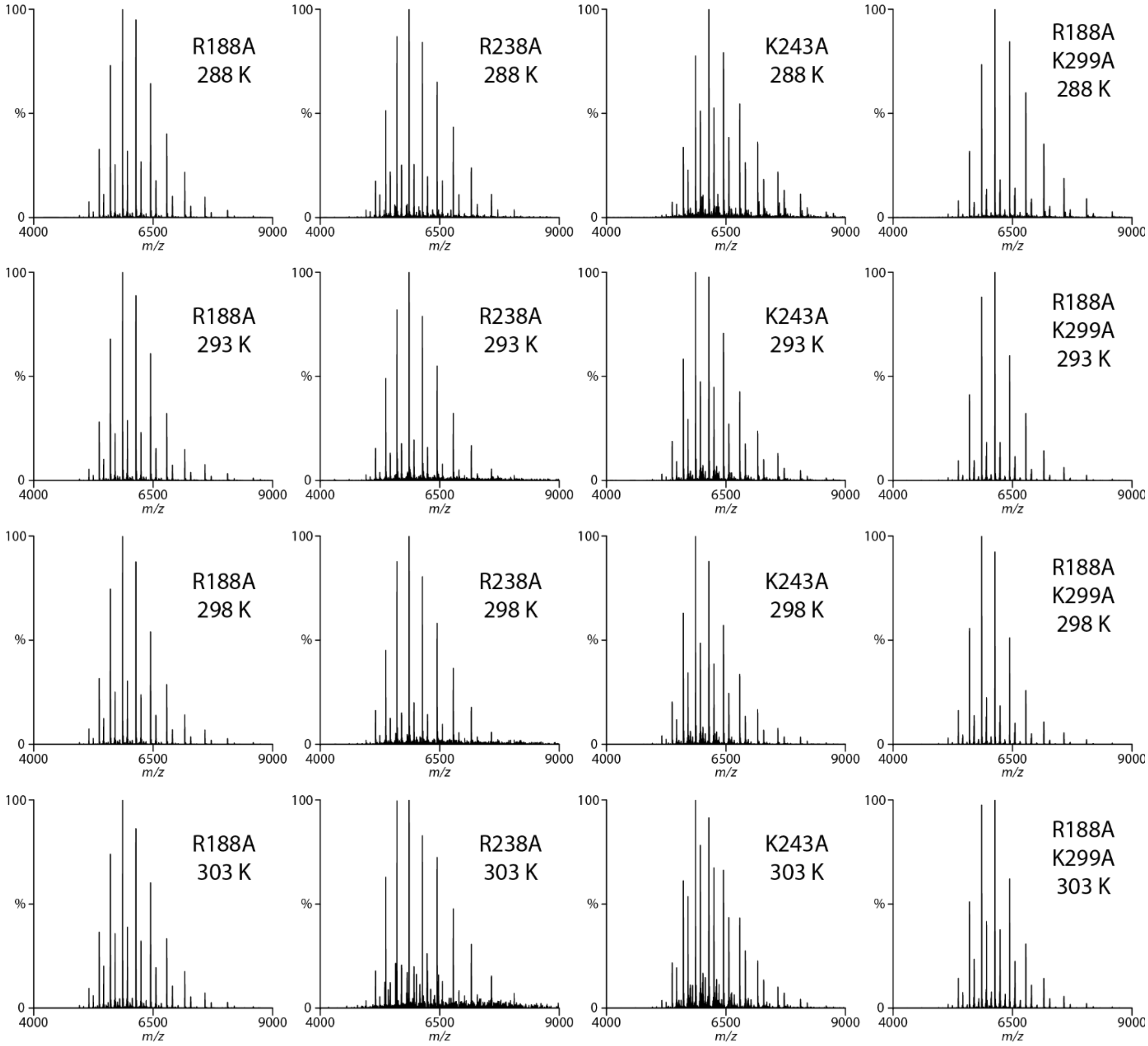


**Supplementary Figure 2**. **Representative native mass spectra MsbA mutants in the presence of 0.8 µM KDL.** The concentration of $MsbA^{R188A}$, $MsbA^{R238A}$, $MsbA^{K243A}$ and $MsbA^{R188A,K299A}$ is 0.56, 0.26, 0.50, and 0.78 µM, respectively. The temperature is provided in the inset.

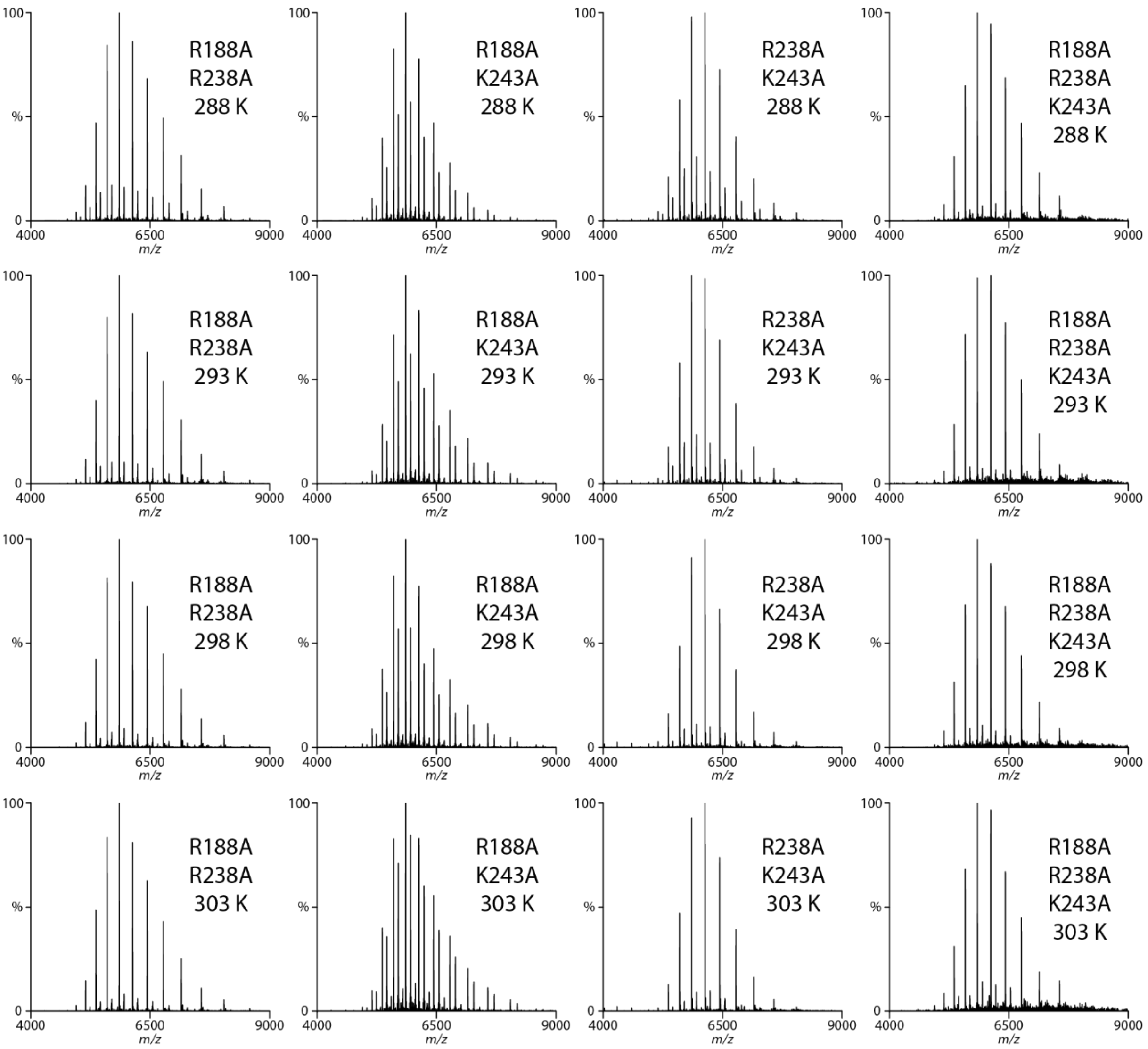


**Supplementary Figure 3**. **Representative native mass spectra MsbA double mutants in the presence of 0.8 µM KDL.** The concentration of $\text{MsbA}^{\text{R188A,R238A}}$, $\text{MsbA}^{\text{R188A,K243A}}$, $\text{MsbA}^{\text{R238A,K243A}}$ and $\text{MsbA}^{\text{R188A,R238A,K243A}}$ is 0.58, 0.27, 0.39, and 0.17 µM, respectively. The temperature is provided in the inset.

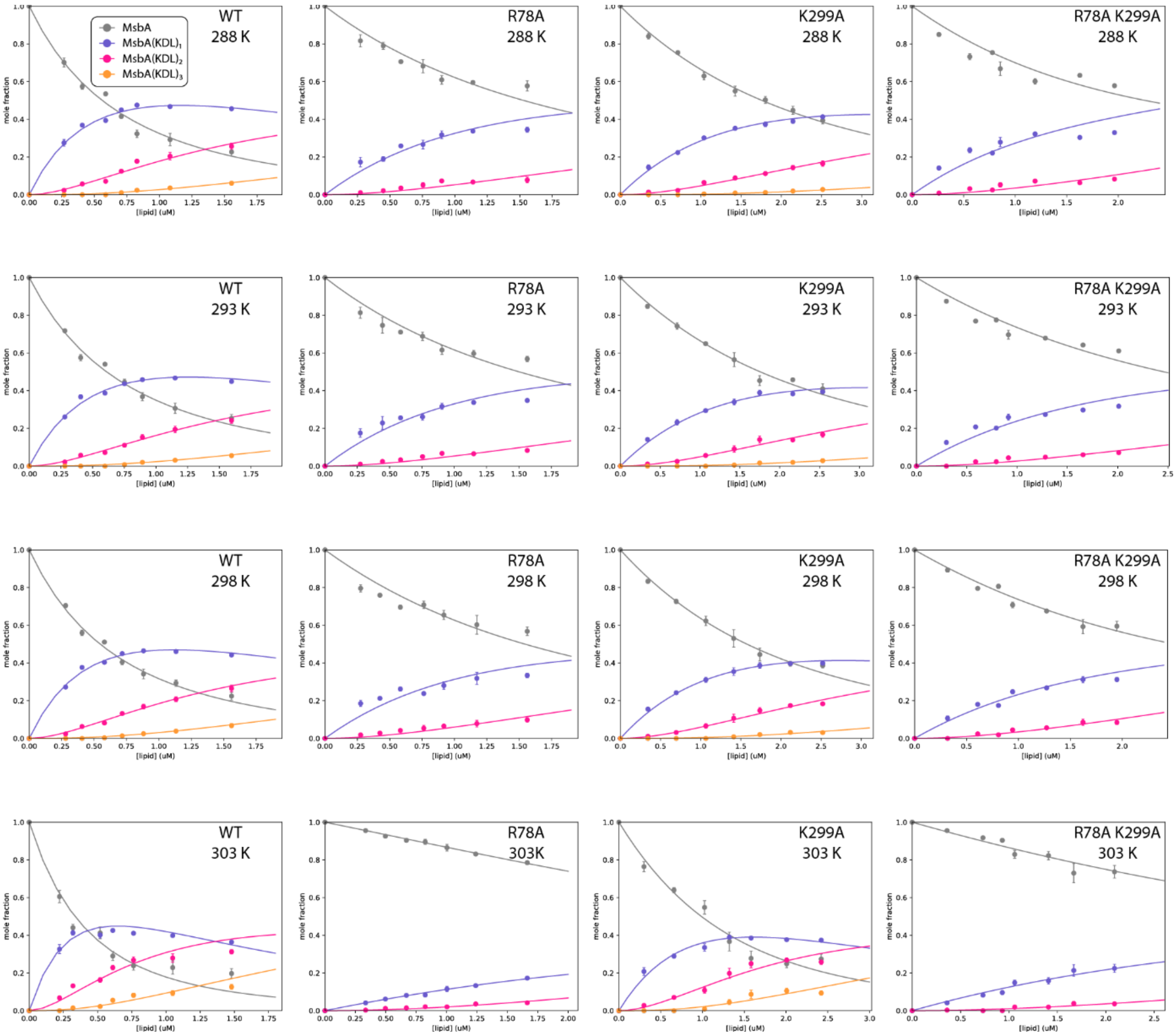


**Supplementary Figure 4**. **Determination of equilibrium dissociation constants ($K_D$) for KDL binding wild-type and mutant MsbA.** Shown are plots of the mole fraction for MsbA and bound to different number of KDL determined from a titration series (dots) and resulting fit from a sequential ligand-binding model (solid lines). The different temperatures and MsbA mutants are labelled. Reported are the mean and standard deviation (*n* = 3).

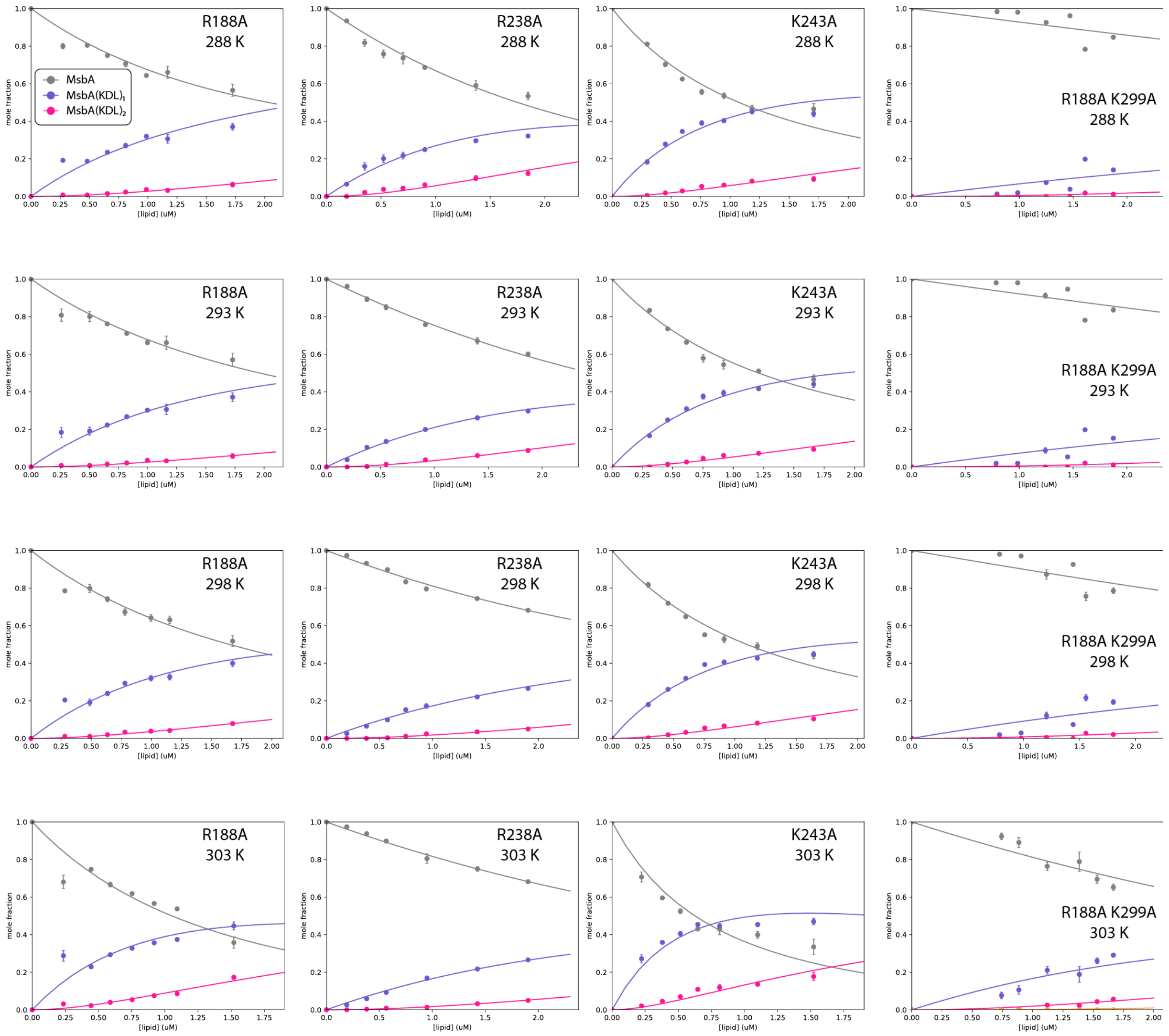


**Supplementary Figure 5. Determination of equilibrium dissociation constants ($K_D$) for MsbA mutants binding KDL.** Shown as described in Supplementary Figure 4.

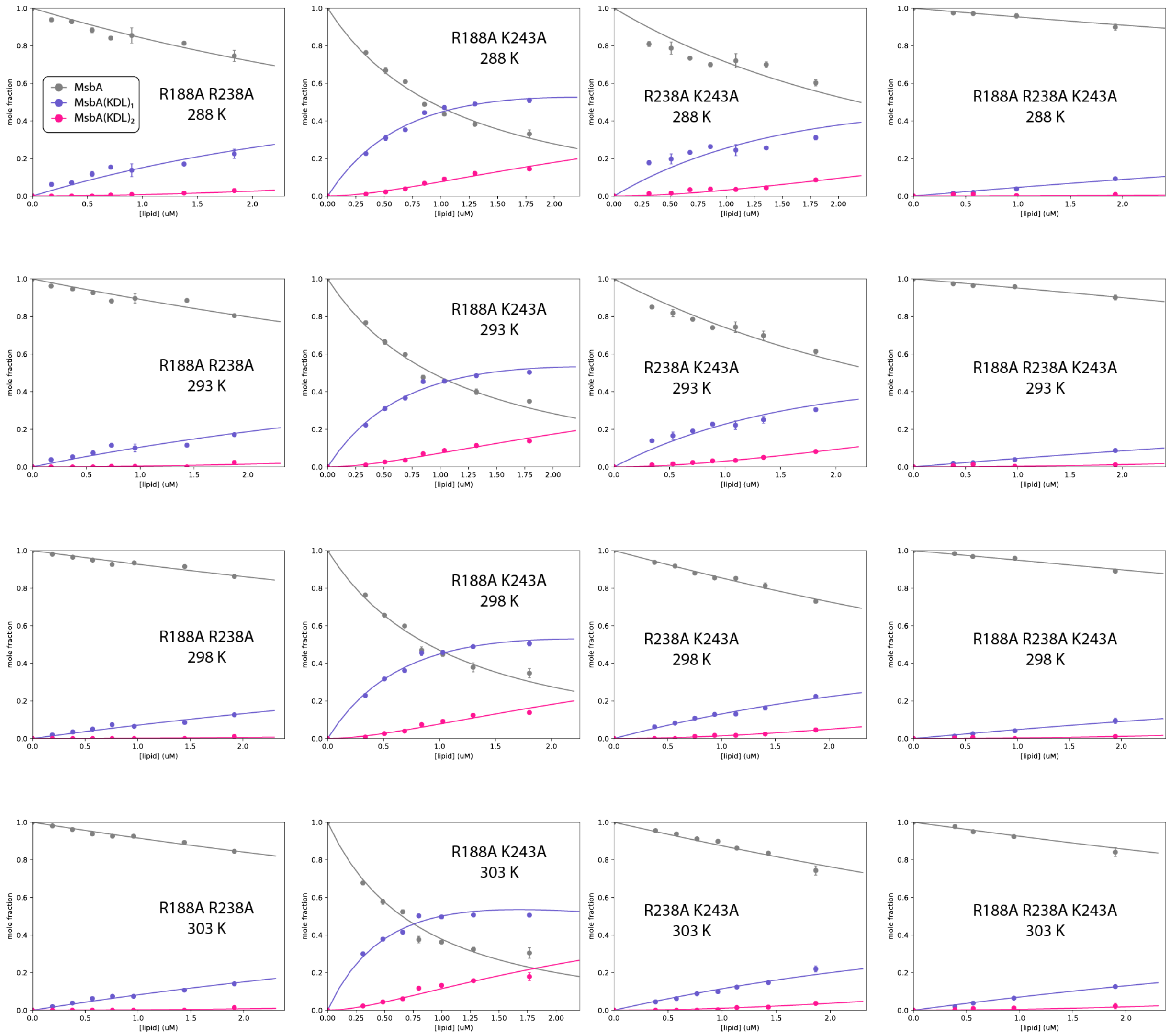


**Supplementary Figure 6. Determination of equilibrium dissociation constants ($K_D$) for MsbA mutants binding KDL.** Shown as described in Supplementary Figure 4.

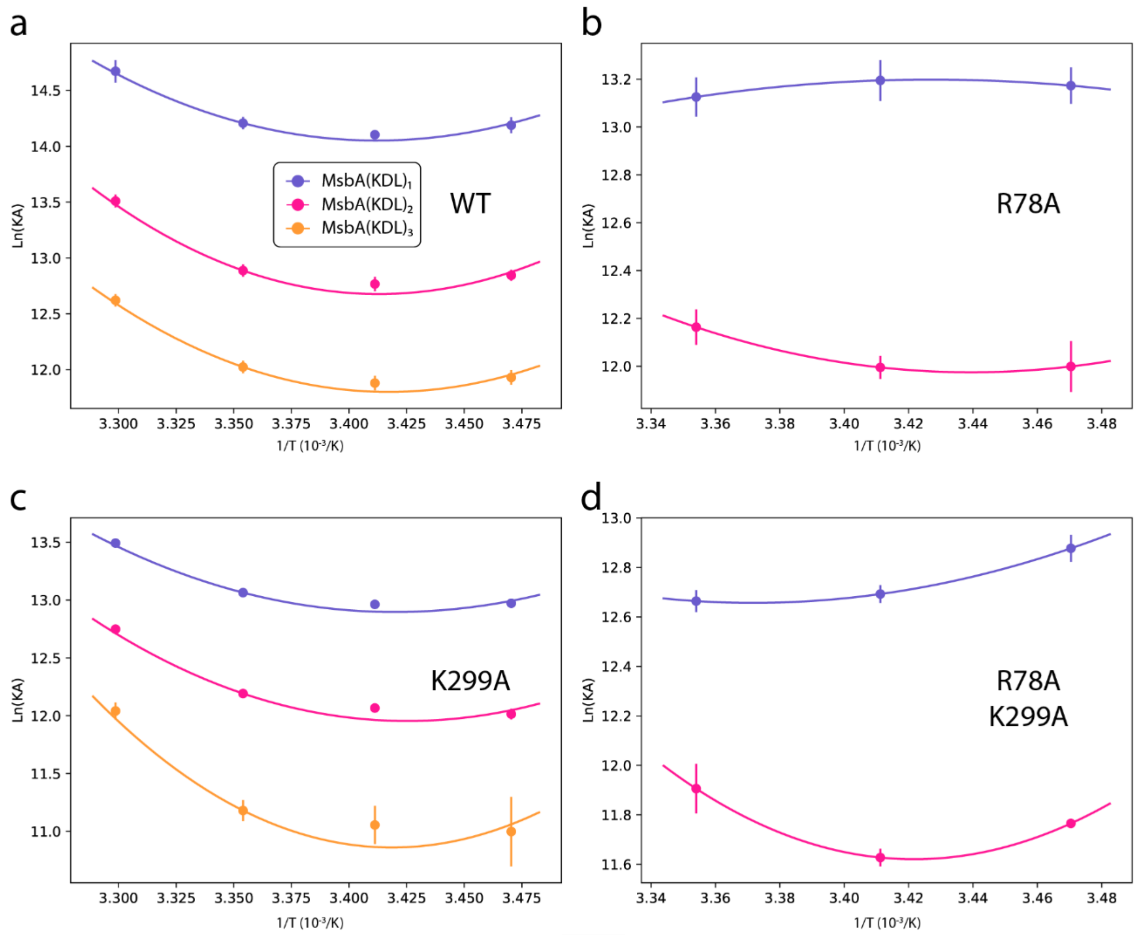


**Supplementary Figure 7. Determination of thermodynamic parameters for KDL binding wild-type and mutant MsbA.** van' t Hoff plots for the transporter binding KDL (dots) and resulting fit from nonlinear Van't Hoff equation (solid lines). Reference temperature ($T_0$) is 298 K Reported are the mean and standard deviation (*n* = 3).

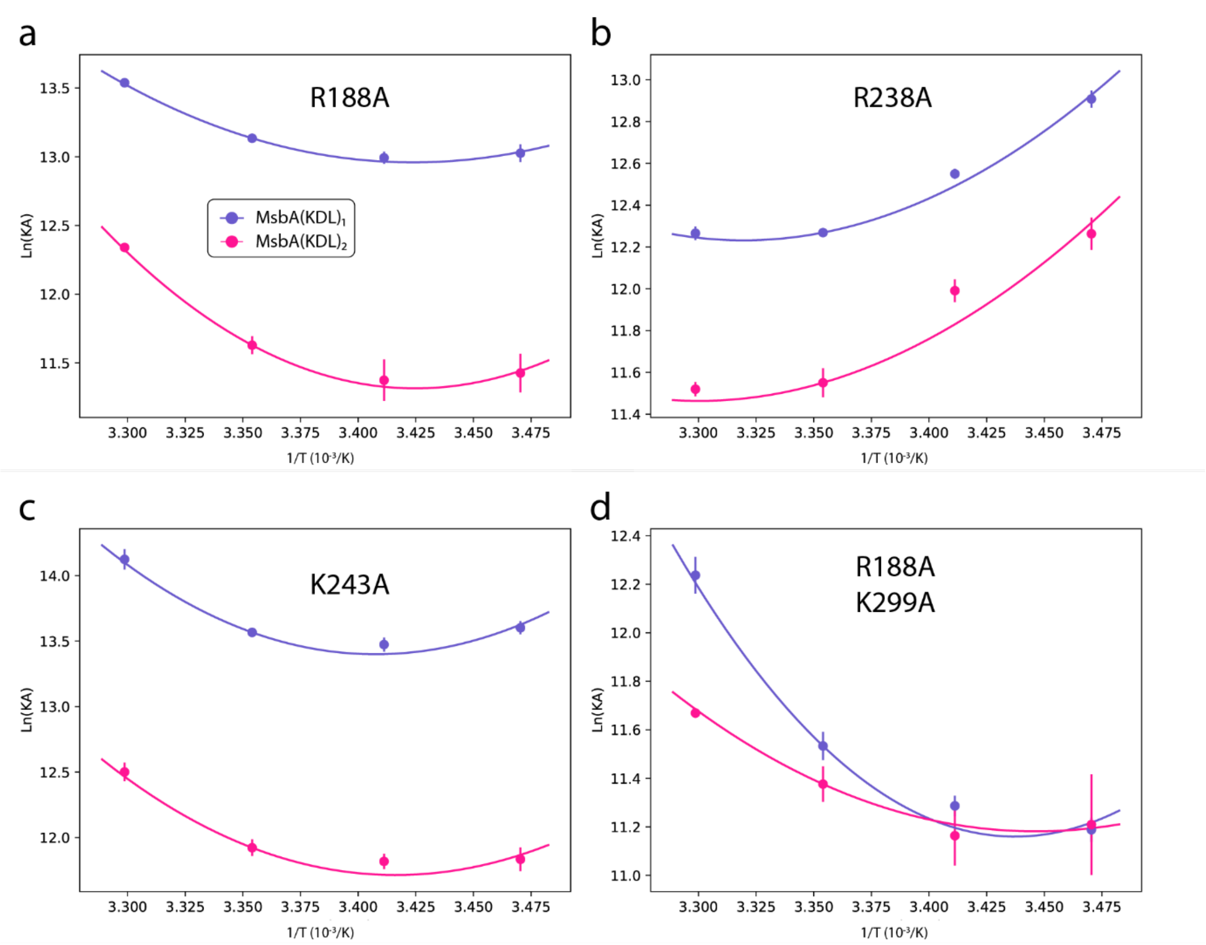


**Supplementary Figure 8. Determination of thermodynamic parameters for KDL binding wild-type and mutant MsbA.** Shown as described in Supplementary Figure 7.

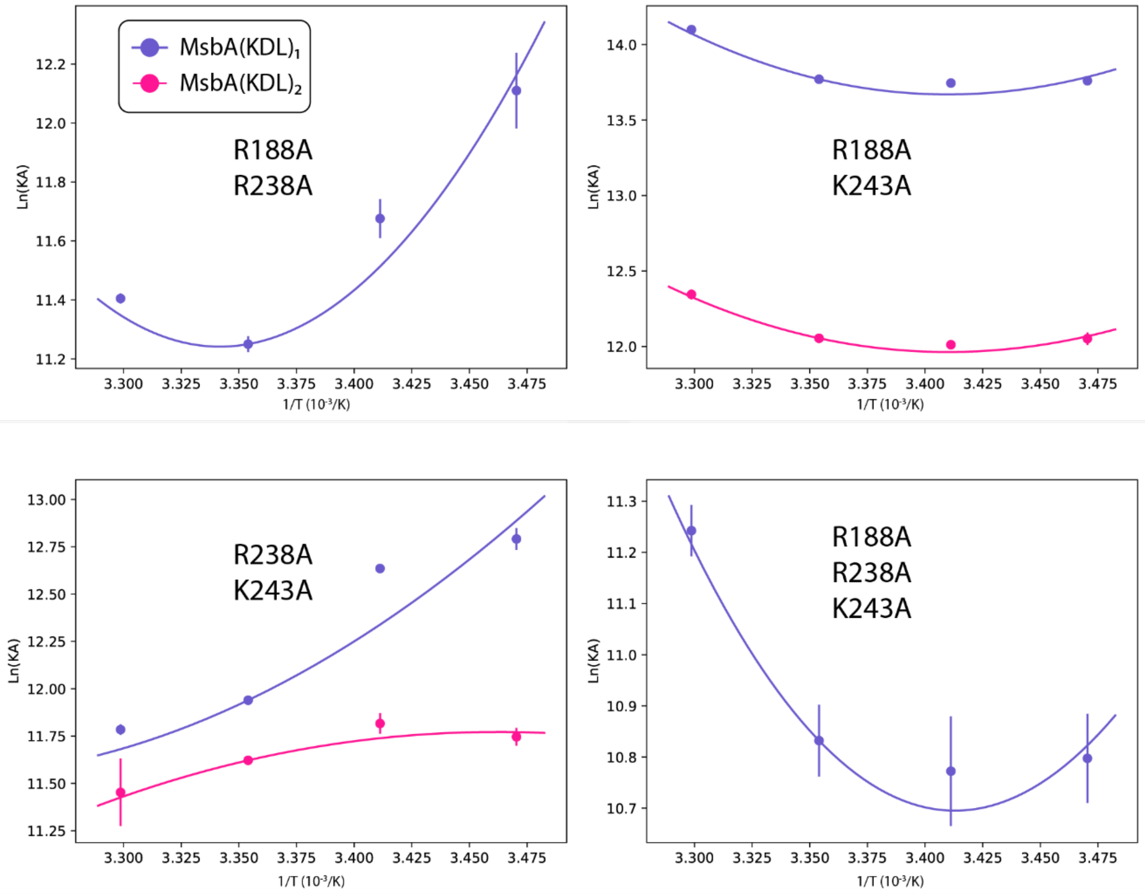


**Supplementary Figure 9. Determination of thermodynamic parameters for KDL binding wild-type and mutant MsbA.** Shown as described in Supplementary Figure 7.

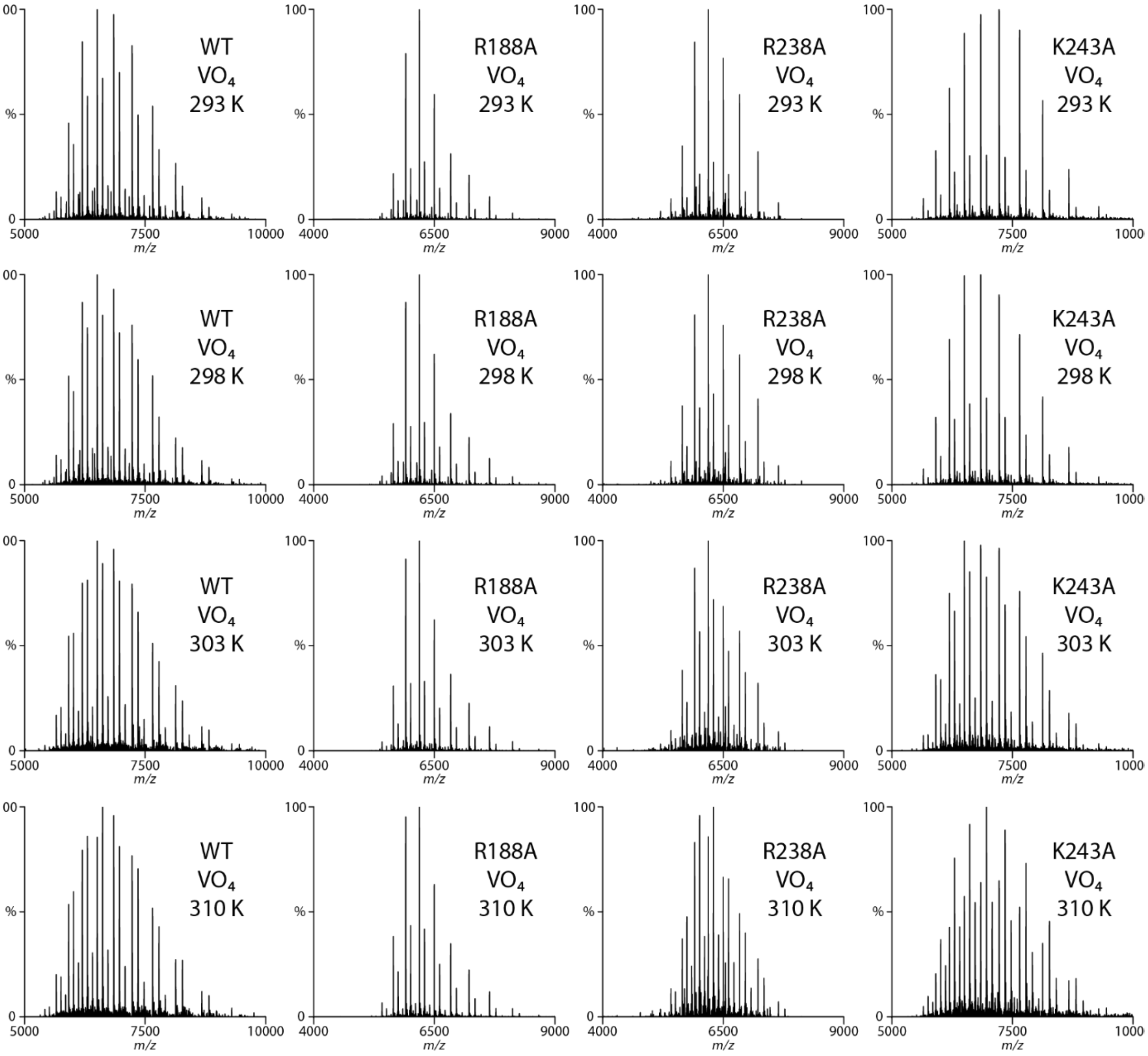


**Supplementary Figure 10**. **Representative native mass spectra wild-type and mutant MsbA trapped with vanadate and ADP in the presence of 0.8 µM KDL.** The concentration of MsbA, $\text{MsbA}^{R188A}$, $\text{MsbA}^{R238A}$ and $\text{MsbA}^{K243A}$ is 0.82, 0.86, 0.78, and 0.57 µM, respectively. The temperature in Celsius is provided in the inset.

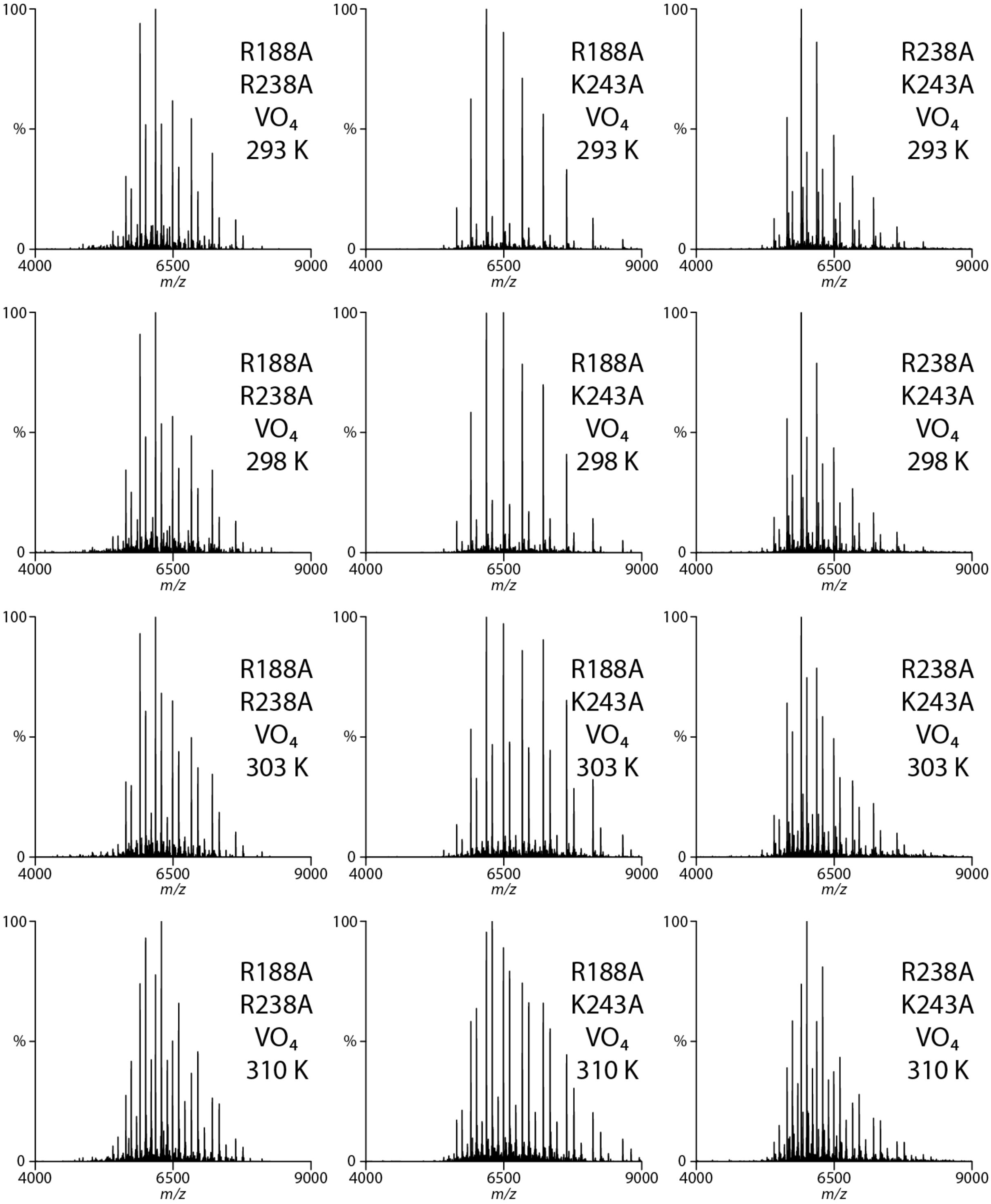


**Supplementary Figure 11**. **Representative native mass spectra MsbA double mutants trapped with vanadate and ADP in the presence of 0.8 µM KDL.** The concentration of $\text{MsbA}^{\text{R188A,R238A}}$, $\text{MsbA}^{\text{R188A,R243A}}$ and $\text{MsbA}^{\text{R238A,K243A}}$ is 0.44, 0.23, and 0.73 µM, respectively. The temperature is provided in the inset.

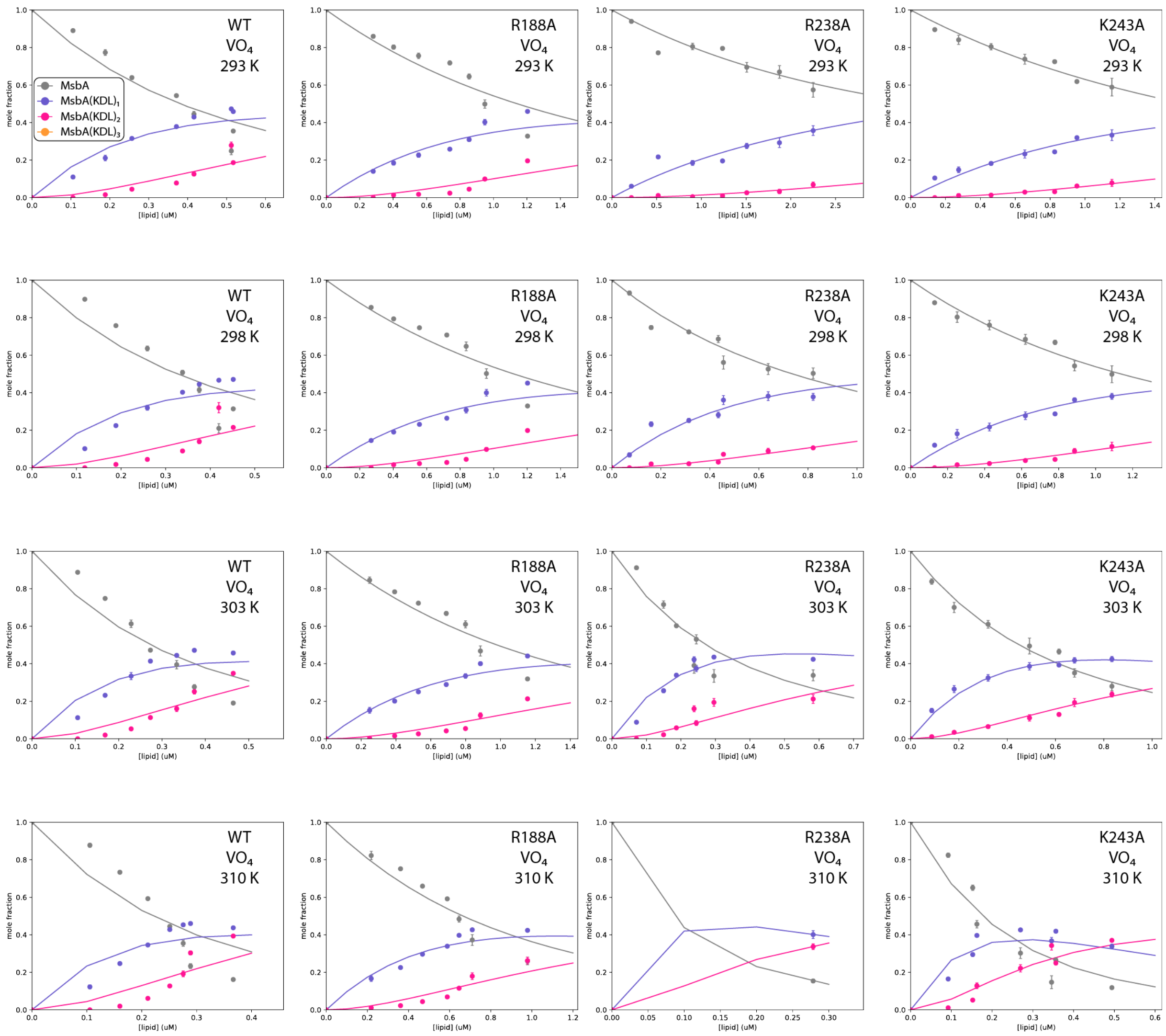


**Supplementary Figure 12. Determination of equilibrium dissociation constants ($K_D$) for KDL binding wild-type and mutant MsbA trapped with ADP and vanadate.** Shown as described in Supplementary Figure 4.

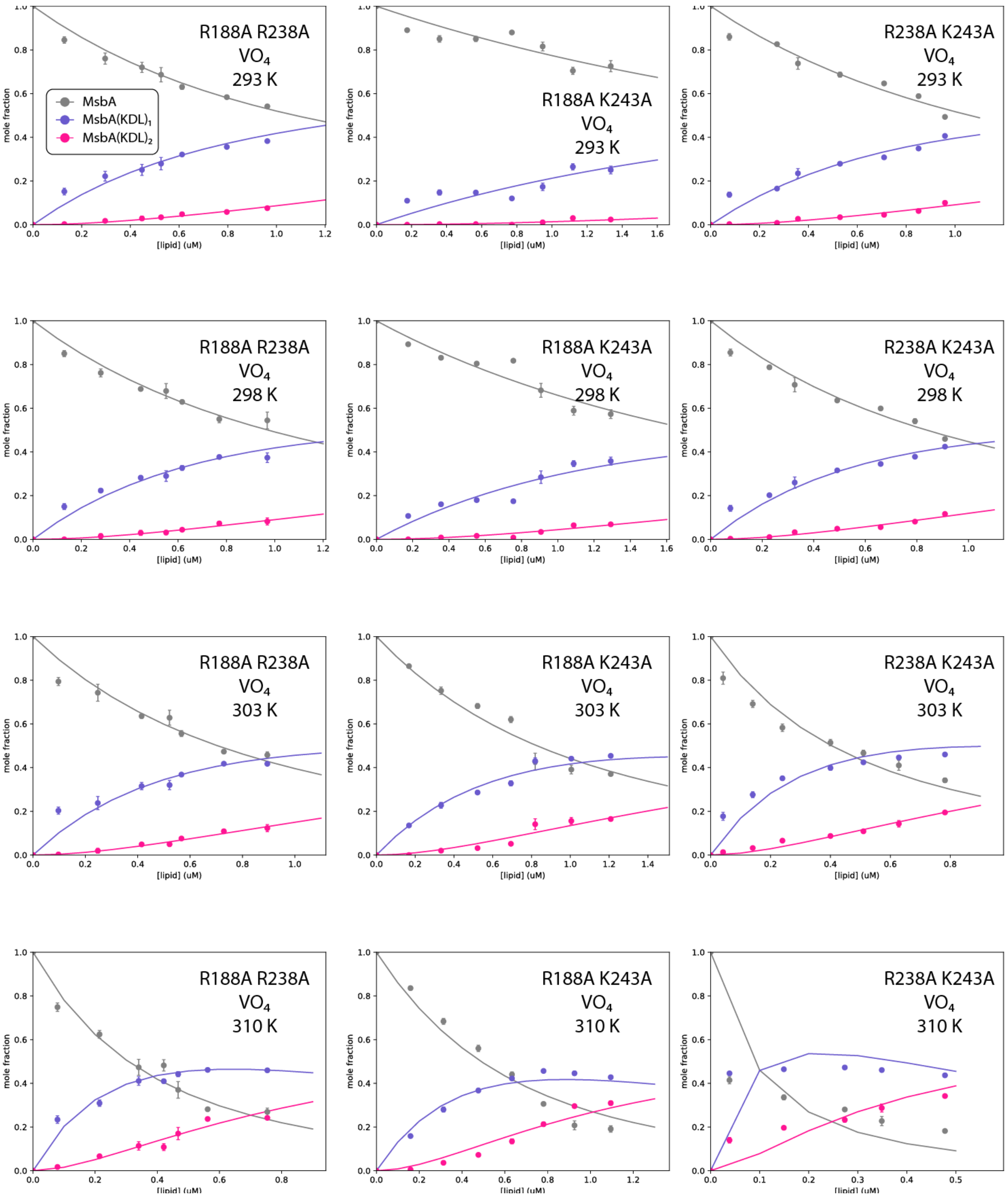


**Supplementary Figure 13. Determination of equilibrium dissociation constants ($K_D$) for KDL binding MsbA double mutants trapped with ADP and vanadate.** Shown as described in Supplementary Figure 4.

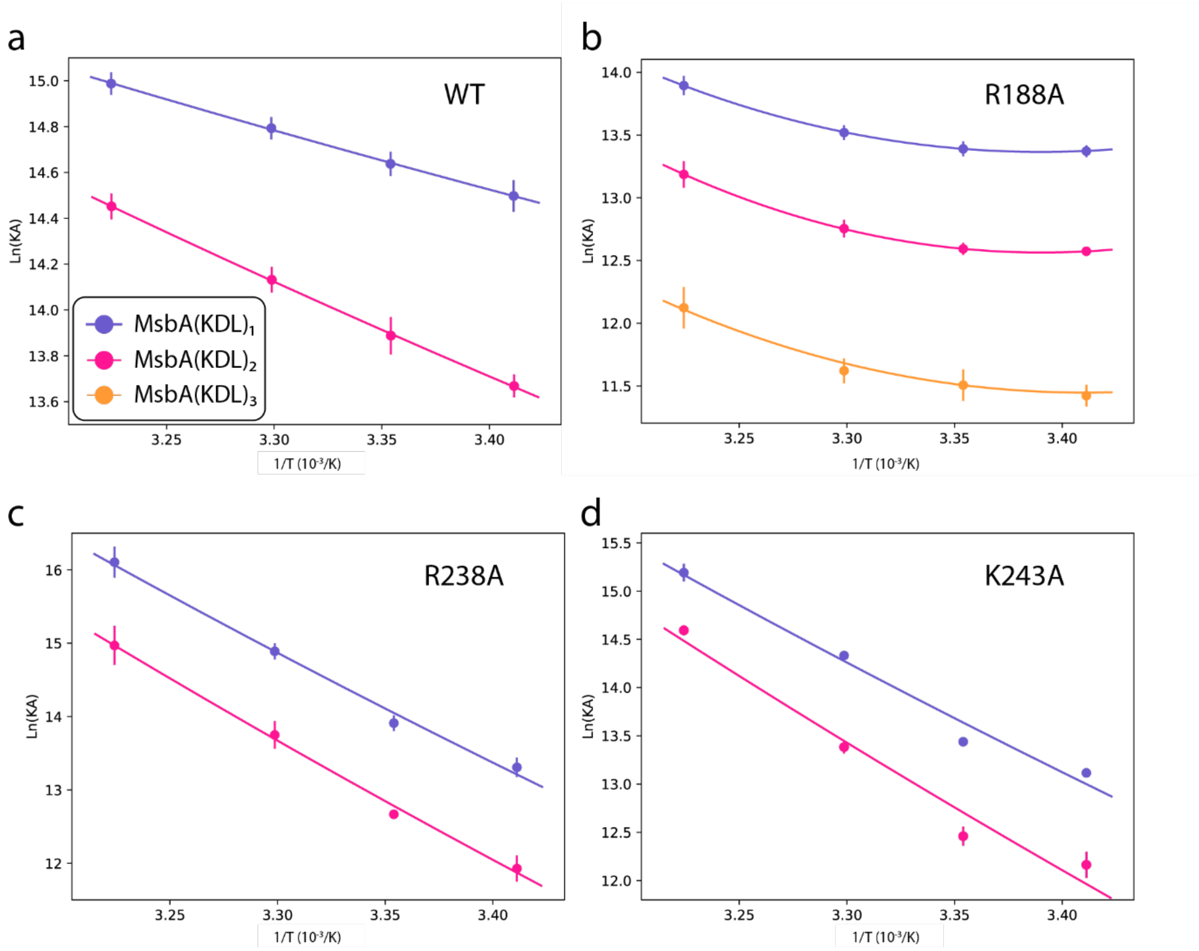


**Supplementary Figure 14. Determination of thermodynamic parameters for KDL binding wild-type and mutant MsbA trapped with ADP and vanadate.** Shown as described in Supplementary Figure 7.

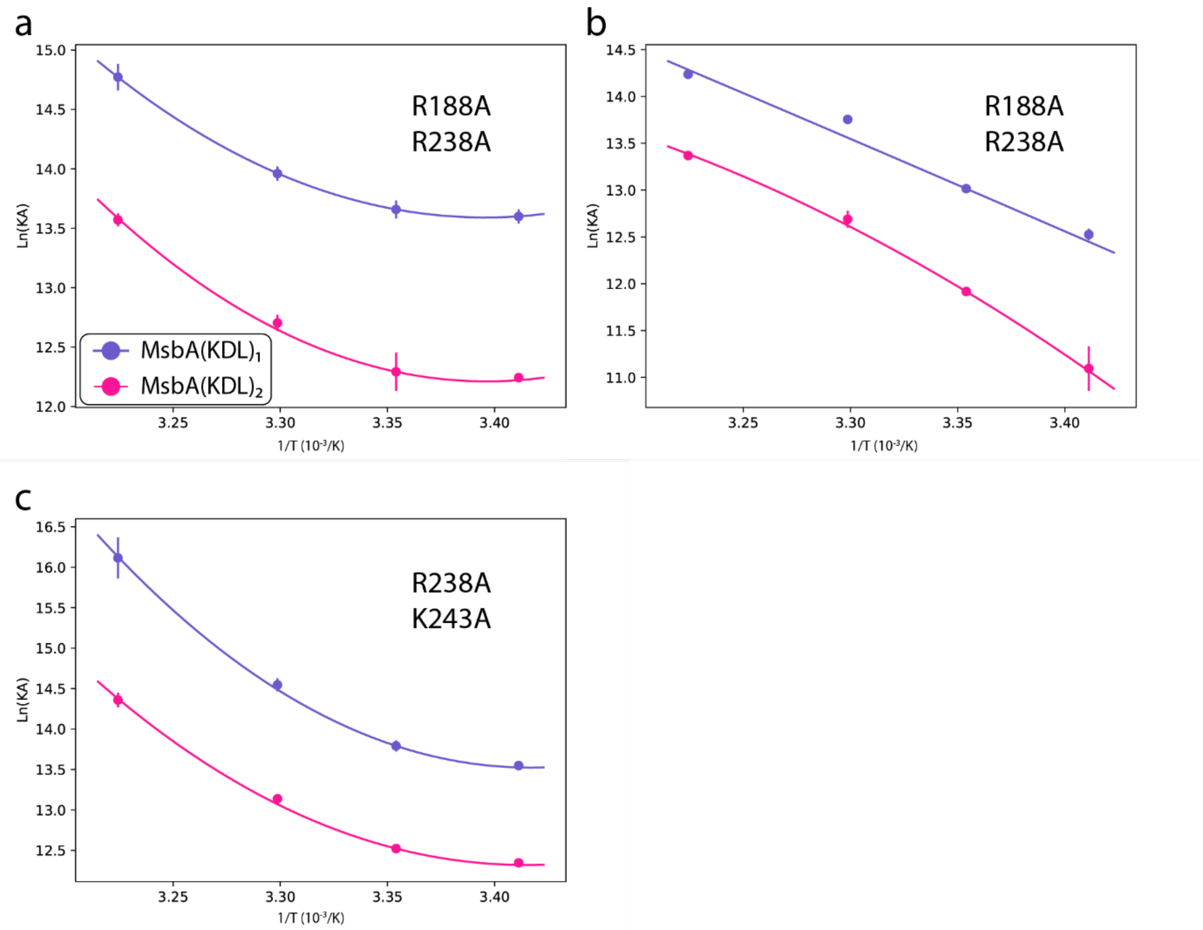


**Supplementary Figure 15. Determination of thermodynamic parameters for KDL binding MsbA double mutants trapped with ADP and vanadate.** Shown as described in Supplementary Figure 7.

**Supplementary table 1. Equilibrium dissociation constants ($K_D$) for KDL binding MsbA at various temperatures.** Reported are the mean and standard deviation (*n* = 3)

| | Temperature(K) | $K_{D1}$(μM) | $K_{D2}$(μM) | $K_{D3}$(μM) | $R^{2*}$ | $X^{2*}$ |
|---|---|---|---|---|---|---|
| **WT** | 288 | 0.69 ± 0.06 | 2.64 ± 0.16 | 6.60 ± 0.54 | 0.99 | 0.01 |
| | 293 | 0.75 ± 0.04 | 2.86 ± 0.23 | 6.94 ± 0.56 | 0.99 | 0.01 |
| | 298 | 0.68 ± 0.05 | 2.53 ± 0.17 | 6.01 ± 0.40 | 0.99 | 0.01 |
| | 303 | 0.43 ± 0.05 | 1.36 ± 0.10 | 3.30 ± 0.23 | 0.96 | 0.06 |
| **R78A** | 288 | 1.91 ± 0.17 | 6.18 ± 0.80 | | 0.98 | 0.04 |
| | 293 | 1.87 ± 0.18 | 6.18 ± 0.37 | | 0.98 | 0.05 |
| | 298 | 2.00 ± 0.20 | 5.23 ± 0.47 | | 0.98 | 0.04 |
| | 303 | 7.68 ± 0.47 | 5.72 ± 0.21 | | 1 | 0 |
| **R188A** | 288 | 2.21 ± 0.17 | 11.01 ± 1.95 | | 0.99 | 0.03 |
| | 293 | 2.28 ± 0.12 | 11.62 ± 2.17 | | 0.99 | 0.03 |
| | 298 | 1.98 ± 0.07 | 8.92 ± 0.73 | | 0.99 | 0.03 |
| | 303 | 1.32 ± 0.05 | 4.37 ± 0.13 | | 0.95 | 0.12 |
| **R238A** | 288 | 2.48 ± 0.12 | 4.74 ± 0.45 | | 0.99 | 0.02 |
| | 293 | 3.55 ± 0.10 | 6.21 ± 0.40 | | 1 | 0.01 |
| | 298 | 4.70 ± 0.07 | 9.66 ± 0.82 | | 1 | 0.01 |
| | 303 | 4.71 ± 0.20 | 9.94 ± 0.40 | | 1 | 0.02 |
| **K243A** | 288 | 1.24 ± 0.07 | 7.28 ± 0.78 | | 0.99 | 0.03 |
| | 293 | 1.41 ± 0.09 | 7.38 ± 0.53 | | 0.99 | 0.02 |
| | 298 | 1.28 ± 0.05 | 6.65 ± 0.50 | | 0.99 | 0.02 |
| | 303 | 0.74 ± 0.07 | 3.73 ± 0.32 | | 0.97 | 0.06 |
| **K299A** | 288 | 2.32 ± 0.06 | 6.07 ± 0.33 | 17.54 ± 6.77 | 1 | 0.01 |
| | 293 | 2.35 ± 0.07 | 5.75 ± 0.07 | 16.03 ± 3.18 | 0.99 | 0.02 |
| | 298 | 2.12 ± 0.09 | 5.07 ± 0.12 | 14.01 ± 1.59 | 0.99 | 0.02 |
| | 303 | 1.38 ± 0.01 | 2.91 ± 0.02 | 5.91 ± 0.55 | 0.97 | 0.05 |
| **R78A K299A** | 288 | 2.56 ± 0.17 | 7.77 ± 0.11 | | 0.98 | 0.06 |
| | 293 | 3.08 ± 0.13 | 8.92 ± 0.39 | | 0.99 | 0.03 |
| | 298 | 3.17 ± 0.17 | 6.78 ± 0.82 | | 0.99 | 0.02 |
| | 303 | 6.91 ± 0.66 | 11.96 ± 2.83 | | 0.99 | 0.03 |
| **R188A R238A** | 288 | 5.55 ± 0.87 | | | 0.99 | 0.02 |
| | 293 | 8.51 ± 0.67 | | | 1 | 0.01 |
| | 298 | 13.01 ± 0.43 | | | 1 | 0 |
| | 303 | 11.14 ± 0.23 | | | 1 | 0 |
| **R188A K243A** | 288 | 1.06 ± 0.02 | 5.84 ± 0.29 | | 0.99 | 0.01 |
| | 293 | 1.07 ± 0.02 | 6.07 ± 0.13 | | 0.99 | 0.01 |
| | 298 | 1.05 ± 0.02 | 5.82 ± 0.06 | | 0.99 | 0.02 |
| | 303 | 0.75 ± 0.01 | 4.35 ± 0.17 | | 0.99 | 0.03 |
| **R188A K299A** | 288 | 13.86 ± 0.85 | 13.83 ± 3.23 | | 0.99 | 0.04 |
| | 293 | 12.55 ± 0.64 | 14.29 ± 2.18 | | 0.99 | 0.03 |
| | 298 | 9.82 ± 0.70 | 11.50 ± 1.04 | | 0.98 | 0.05 |
| | 303 | 4.86 ± 0.44 | 8.56 ± 0.20 | | 0.99 | 0.04 |
| **R238A K243A** | 288 | 2.79 ± 0.20 | 7.93 ± 0.47 | | 0.98 | 0.06 |
| | 293 | 3.26 ± 0.09 | 7.39 ± 0.50 | | 0.99 | 0.03 |
| | 298 | 6.53 ± 0.2 | 8.97 ± 0.15 | | 1 | 0 |
| | 303 | 7.63 ± 0.26 | 10.78 ± 2.19 | | 1 | 0.01 |
| **R188A R238A K243A** | 288 | 20.53 ± 2.25 | | | 1 | 0 |
| | 293 | 21.09 ± 2.84 | | | 1 | 0 |
| | 298 | 19.80 ± 1.71 | | | 1 | 0 |
| | 303 | 13.12 ± 0.81 | | | 1 | 0 |

*These values represent the replicates with the poorest fits.

**Supplementary table 2. Thermodynamic signatures of KDL interacting with wild-type and mutant MsbA.** Reported are the mean with standard deviation (*n* = 3), and the subscript denotes the $n^{th}$ KDL binding event.

| | T (K) | $\Delta G_1$ (kJ/mol) | $\Delta H_1$ (kJ/mol) | $-T\Delta S_1$ (kJ/mol) | $\Delta Cp_1$ (kJ/mol/K) | $\Delta G_2$ (kJ/mol) | $\Delta H_2$ (kJ/mol) | $-T\Delta S_2$ (kJ/mol) | $\Delta Cp_2$ (kJ/mol/K) | $\Delta G_3$ (kJ/mol) | $\Delta H_3$ (kJ/mol) | $-T\Delta S_3$ (kJ/mol) | $\Delta Cp_3$ (kJ/mol/K) |
|---|---|---|---|---|---|---|---|---|---|---|---|---|---|
| **WT** | 288 | -34.0 ± 0.2 | -36.0 ± 11.6 | 2.0 ± 11.5 | 8.0 ± 1.5 | -30.8 ± 0.1 | -43.2 ± 7.3 | 12.4 ± 7.3 | 10.1 ± 0.9 | -28.6 ± 0.2 | -36.6 ± 8.1 | 8.0 ± 8.0 | 9.4 ± 0.9 |
| | 293 | -34.4 ± 0.1 | 4.3 ± 5.4 | -38.6 ± 5.4 | 7.4 ± 1.5 | -31.1 ± 0.2 | 7.9 ± 2.7 | -39.0 ± 2.7 | 9.1 ± 0.8 | -29.0 ± 0.2 | 10.9 ± 3.3 | -39.8 ± 3.3 | 8.5 ± 0.6 |
| | 298 | -35.2 ± 0.2 | 45.0 ± 5.8 | -80.2 ± 5.9 | 8.9 ± 1.4 | -32.0 ± 0.2 | 59.9 ± 2.3 | -91.8 ± 2.2 | 11.7 ± 1.2 | -29.8 ± 0.2 | 59.1 ± 1.8 | -88.9 ± 1.7 | 10.7 ± 1.5 |
| | 303 | -37.0 ± 0.3 | 86.1 ± 12.1 | -123.0 ± 12.4 | 8.3 ± 1.5 | -34.1 ± 0.2 | 112.3 ± 7.1 | -146.4 ± 7.1 | 10.6 ± 1.0 | -31.8 ± 0.2 | 107.7 ± 7.1 | -139.5 ± 7.1 | 9.8 ± 1.1 |
| **R78A** | 288 | -31.6 ± 0.2 | 9.6 ± 4.5 | -41.2 ± 4.7 | -2.6 ± 1.1 | -28.8 ± 0.3 | -13.0 ± 19.1 | -15.8 ± 18.8 | 5.0 ± 1.7 | | | | |
| | 293 | -32.2 ± 0.3 | -3.5 ± 5.1 | -28.6 ± 5.1 | -2.6 ± 1.1 | -29.2 ± 0.1 | 12.0 ± 11.1 | -41.3 ± 11.1 | 5.0 ± 1.7 | | | | |
| | 298 | -32.5 ± 0.2 | -16.7 ± 9.6 | -15.9 ± 9.6 | -2.6 ± 1.1 | -30.2 ± 0.2 | 37.1 ± 5.9 | -67.2 ± 6.1 | 5.0 ± 1.7 | | | | |
| **R188A** | 288 | -31.2 ± 0.2 | -23.0 ± 8.2 | -8.3 ± 8.1 | 6.4 ± 1.1 | -27.4 ± 0.4 | -39.1 ± 0.5 | 11.8 ± 0.7 | 11.2 ± 1.0 | | | | |
| | 293 | -31.7 ± 0.1 | 9.4 ± 3.6 | -41.1 ± 3.5 | 6.1 ± 1.2 | -27.7 ± 0.5 | 17.4 ± 4.6 | -45.1 ± 4.2 | 10.7 ± 1.7 | | | | |
| | 298 | -32.6 ± 0.1 | 42.1 ± 3.7 | -74.6 ± 3.7 | 6.9 ± 0.9 | -28.8 ± 0.2 | 74.4 ± 8.8 | -103.2 ± 8.6 | 12.1 ± 0.7 | | | | |
| | 303 | -34.1 ± 0.1 | 74.9 ± 8.3 | -109.0 ± 8.3 | 6.6 ± 1.0 | -31.1 ± 0.1 | 131.6 ± 12.8 | -162.7 ± 12.8 | 11.5 ± 0.7 | | | | |
| **R238A** | 288 | -30.9 ± 0.1 | -66.8 ± 3.4 | 35.9 ± 3.3 | 4.8 ± 0.3 | -29.4 ± 0.2 | -57.9 ± 17.9 | 28.6 ± 17.8 | 2.8 ± 2.4 | | | | |
| | 293 | -30.6 ± 0.1 | -42.6 ± 4.0 | 12.1 ± 3.9 | 4.1 ± 0.3 | -29.2 ± 0.2 | -43.2 ± 6.5 | 13.9 ± 6.5 | 0.9 ± 2.5 | | | | |
| | 298 | -30.4 ± 0.1 | -17.9 ± 4.8 | -12.6 ± 4.8 | 5.8 ± 0.3 | -28.6 ± 0.2 | -26.8 ± 7.2 | -1.9 ± 7.0 | 5.5 ± 2.4 | | | | |
| | 303 | -30.9 ± 0.1 | 7.2 ± 5.8 | -38.1 ± 5.9 | 5.1 ± 0.3 | -29.0 ± 0.1 | -9.5 ± 18.6 | -19.5 ± 18.7 | 3.7 ± 2.4 | | | | |
| **K243A** | 288 | -32.6 ± 0.1 | -48.5 ± 7.3 | 15.9 ± 7.2 | 9.9 ± 0.9 | -28.4 ± 0.3 | -31.6 ± 7.3 | 3.3 ± 7.0 | 8.5 ± 0.9 | | | | |
| | 293 | -32.8 ± 0.2 | 1.5 ± 3.5 | -34.4 ± 3.5 | 9.1 ± 0.8 | -28.8 ± 0.2 | 11.7 ± 3.1 | -40.5 ± 3.0 | 7.3 ± 1.2 | | | | |
| | 298 | -33.6 ± 0.1 | 52.2 ± 3.9 | -85.9 ± 4.0 | 11.2 ± 1.2 | -29.6 ± 0.2 | 56.0 ± 2.0 | -85.5 ± 2.1 | 10.3 ± 0.6 | | | | |
| | 303 | -35.6 ± 0.2 | 103.3 ± 8.0 | -138.9 ± 8.2 | 10.3 ± 1.0 | -31.5 ± 0.2 | 100.8 ± 5.6 | -132.3 ± 5.8 | 9.1 ± 0.8 | | | | |
| **K299A** | 288 | -31.1 ± 0.1 | -22.2 ± 9.5 | -8.9 ± 9.4 | 6.4 ± 1.1 | -28.8 ± 0.1 | -18.8 ± 9.6 | -10.0 ± 9.5 | 7.3 ± 1.0 | -26.4 ± 0.9 | -35.6 ± 71.5 | 9.3 ± 70.7 | 11.4 ± 7.1 |
| | 293 | -31.6 ± 0.1 | 9.9 ± 3.8 | -41.5 ± 3.9 | 5.6 ± 1.1 | -29.4 ± 0.1 | 18.0 ± 4.6 | -47.4 ± 4.6 | 6.0 ± 1.0 | -27.0 ± 0.5 | 22.7 ± 37.2 | -49.6 ± 37.5 | 9.2 ± 8.6 |
| | 298 | -32.4 ± 0.1 | 42.7 ± 2.1 | -75.1 ± 1.9 | 7.4 ± 1.2 | -30.2 ± 0.1 | 55.9 ± 1.7 | -86.1 ± 1.6 | 9.0 ± 1.1 | -27.7 ± 0.3 | 82.9 ± 7.8 | -110.6 ± 7.9 | 14.7 ± 5.1 |
| | 303 | -34.0 ± 0.1 | 75.8 ± 7.7 | -109.8 ± 7.7 | 6.7 ± 1.2 | -32.1 ± 0.1 | 94.3 ± 6.3 | -126.4 ± 6.3 | 7.8 ± 1.1 | -30.4 ± 0.2 | 144.1 ± 30.5 | -174.5 ± 30.3 | 12.5 ± 6.4 |
| **R78A K299A** | 288 | -30.9 ± 0.2 | -36.8 ± 5.6 | 5.9 ± 5.4 | 4.4 ± 0.8 | -28.2 ± 0.1 | -49.1 ± 5.7 | 20.9 ± 5.7 | 12.0 ± 1.4 | | | | |
| | 293 | -30.9 ± 0.1 | -15.0 ± 2.8 | -16.0 ± 2.8 | 4.4 ± 0.8 | -28.3 ± 0.1 | 10.7 ± 9.6 | -39.1 ± 9.7 | 12.0 ± 1.4 | | | | |
| | 298 | -31.4 ± 0.1 | 6.8 ± 3.8 | -38.2 ± 3.9 | 4.4 ± 0.8 | -29.5 ± 0.3 | 70.6 ± 15.6 | -100.1 ± 15.9 | 12.0 ± 1.4 | | | | |
| **R188A R238A** | 288 | -29.0 ± 0.4 | -93.3 ± 15.2 | 64.3 ± 14.8 | 7.8 ± 1.3 | | | | | | | | |
| | 293 | -28.5 ± 0.2 | -53.3 ± 9.2 | 24.8 ± 9.1 | 5.9 ± 1.6 | | | | | | | | |
| | 298 | -27.9 ± 0.1 | -11.6 ± 4.7 | -16.3 ± 4.6 | 10.6 ± 1.0 | | | | | | | | |
| | 303 | -28.8 ± 0.1 | 30.9 ± 5.9 | -59.7 ± 5.9 | 8.7 ± 1.2 | | | | | | | | |
| **R188A K243A** | 288 | -33 ± 0.1 | -20.3 ± 6.0 | -12.6 ± 6.0 | 4.9 ± 0.6 | -28.9 ± 0.1 | -21.3 ± 3.4 | -7.6 ± 3.3 | 4.8 ± 0.5 | | | | |
| | 293 | -33.5 ± 0.1 | 4.5 ± 3.2 | -38.0 ± 3.2 | 4.0 ± 0.6 | -29.3 ± 0.1 | 2.7 ± 3.4 | -32.0 ± 3.4 | 4.2 ± 0.4 | | | | |
| | 298 | -34.1 ± 0.1 | 30.1 ± 1.9 | -64.3 ± 1.9 | 6.2 ± 0.7 | -29.9 ± 0.1 | 27.3 ± 5.0 | -57.1 ± 5.0 | 5.6 ± 0.6 | | | | |
| | 303 | -35.5 ± 0.1 | 56.1 ± 4.1 | -91.7 ± 4.1 | 5.3 ± 0.6 | -31.1 ± 0.1 | 52.0 ± 7.3 | -83.2 ± 7.4 | 5.0 ± 0.5 | | | | |
| **R188A K299A** | 288 | -26.8 ± 0.1 | -15.4 ± 16.2 | -11.4 ± 16.0 | 8.9 ± 2.2 | -26.9 ± 0.6 | -15.7 ± 36.0 | -11.1 ± 35.4 | 5.2 ± 3.8 | | | | |
| | 293 | -27.5 ± 0.1 | 29.5 ± 5.2 | -57.0 ± 5.2 | 7.8 ± 1.3 | -27.2 ± 0.4 | 10.1 ± 18.9 | -37.3 ± 18.9 | 5.8 ± 6.0 | | | | |
| | 298 | -28.6 ± 0.2 | 75.2 ± 7.6 | -103.8 ± 7.4 | 10.4 ± 3.5 | -28.2 ± 0.2 | 35.4 ± 6.4 | -63.6 ± 6.3 | 4.4 ± 1.4 | | | | |
| | 303 | -30.8 ± 0.2 | 121.4 ± 19.9 | -152.2 ± 20.0 | 9.4 ± 2.6 | -29.4 ± 0.1 | 60.5 ± 14.2 | -89.9 ± 14.2 | 5.0 ± 2.8 | | | | |
| **R238A K243A** | 288 | -30.7 ± 0.2 | -42.4 ± 8.3 | 11.8 ± 8.1 | -1.2 ± 0.5 | -28.1 ± 0.1 | 13.2 ± 9.6 | -41.4 ± 9.6 | -3.8 ± 2.4 | | | | |
| | 293 | -30.8 ± 0.1 | -46.9 ± 6.0 | 16.1 ± 6.0 | -4.7 ± 0.3 | -28.8 ± 0.2 | -5.4 ± 4.5 | -23.4 ± 4.5 | -4.7 ± 1.6 | | | | |
| | 298 | -29.6 ± 0.1 | -48.4 ± 4.0 | 18.8 ± 4.1 | 3.8 ± 0.8 | -28.8 ± 0.1 | -23.3 ± 17.2 | -5.5 ± 17.2 | -2.5 ± 3.8 | | | | |
| | 303 | -29.7 ± 0.1 | -48.3 ± 3.4 | 18.6 ± 3.4 | 0.4 ± 0.6 | -28.9 ± 0.6 | -40.7 ± 31 | 11.9 ± 31.5 | -3.4 ± 2.9 | | | | |
| **R188A R238A K243A** | 288 | -25.9 ± 0.3 | -25.2 ± 5.7 | -0.7 ± 5.4 | 6.2 ± 0.3 | | | | | | | | |
| | 293 | -26.3 ± 0.3 | 6.5 ± 3.7 | -32.7 ± 3.4 | 5.3 ± 1.4 | | | | | | | | |
| | 298 | -26.9 ± 0.2 | 38.9 ± 0.6 | -65.7 ± 0.7 | 7.6 ± 2.5 | | | | | | | | |
| | 303 | -28.3 ± 0.2 | 71.7 ± 3.3 | -100 ± 3.3 | 6.7 ± 0.9 | | | | | | | | |

**Supplementary Table 3. Double mutant cycle analysis of the first KDL binding to wild-type and mutant MsbA.** The ΔΔ values mutant relative to the wild-type protein. Reported are the mean (*n* = 3).

| | | Temperature (K) | ΔΔG (kJ/mol) | ΔΔH (kJ/mol) | Δ(-TΔS) (kJ/mol) | $\Delta\Delta G_{int}$ (kJ/mol) | $\Delta\Delta H_{int}$ (kJ/mol) | $\Delta(-\Delta TS)_{int}$ (kJ/mol) |
|---|---|---|---|---|---|---|---|---|
| **R78A** | | 288 | 2.4 ± 0.2 | 45.6 ± 12.5 | -43.2 ± 12.4 | | | |
| | | 293 | 2.2 ± 0.2 | -7.8 ± 7.5 | 10.0 ± 7.5 | | | |
| | | 298 | 2.7 ± 0.2 | -61.7 ± 11.3 | 64.4 ± 11.3 | | | |
| **R188A** | | 288 | 2.8 ± 0.2 | 13.0 ± 14.2 | -10.2 ± 14.0 | | | |
| | | 293 | 2.7 ± 0.1 | 5.2 ± 6.5 | -2.5 ± 6.5 | | | |
| | | 298 | 2.7 ± 0.2 | -3.0 ± 6.9 | 5.6 ± 7.0 | | | |
| | | 303 | 2.9 ± 0.4 | -11.2 ± 14.7 | 14.1 ± 14.9 | | | |
| **R238A** | | 288 | 3.1 ± 0.2 | -30.9 ± 12.1 | 33.9 ± 11.9 | | | |
| | | 293 | 3.8 ± 0.1 | -46.9 ± 6.7 | 50.7 ± 6.7 | | | |
| | | 298 | 4.8 ± 0.1 | -62.9 ± 7.5 | 67.7 ± 7.6 | | | |
| | | 303 | 6.1 ± 0.4 | -78.8 ± 13.5 | 84.9 ± 13.7 | | | |
| **K243A** | | 288 | 1.4 ± 0.2 | -12.5 ± 13.7 | 13.9 ± 13.5 | | | |
| | | 293 | 1.5 ± 0.2 | -2.7 ± 6.5 | 4.3 ± 6.5 | | | |
| | | 298 | 1.6 ± 0.2 | 7.2 ± 7.0 | -5.6 ± 7.1 | | | |
| | | 303 | 1.4 ± 0.4 | 17.3 ± 14.5 | -15.9 ± 14.8 | | | |
| **K299A** | | 288 | 2.9 ± 0.2 | 13.8 ± 15.1 | -10.9 ± 14.8 | | | |
| | | 293 | 2.8 ± 0.1 | 5.7 ± 6.6 | -2.9 ± 6.7 | | | |
| | | 298 | 2.8 ± 0.2 | -2.4 ± 6.1 | 5.2 ± 6.2 | | | |
| | | 303 | 3.0 ± 0.4 | -10.3 ± 14.3 | 13.3 ± 14.6 | | | |
| **R78A** | **K299A** | 288 | 3.2 ± 0.2 | -0.8 ± 12.9 | 4.0 ± 12.6 | 2.2 ± 0.5 | 60.2 ± 23.4 | -58.0 ± 23.1 |
| | | 293 | 3.4 ± 0.1 | -19.2 ± 6.1 | 22.7 ± 6.1 | 1.6 ± 0.4 | 17.1 ± 11.6 | -15.6 ± 11.8 |
| | | 298 | 3.8 ± 0.2 | -38.2 ± 6.9 | 42.0 ± 7.1 | 1.7 ± 0.4 | -25.8 ± 14.6 | 27.5 ± 14.7 |
| **R188A** | **R238A** | 288 | 5.0 ± 0.5 | -57.4 ± 19.1 | 62.4 ± 18.7 | 0.9 ± 0.6 | 39.5 ± 26.7 | -38.7 ± 26.2 |
| | | 293 | 5.9 ± 0.2 | -57.5 ± 10.7 | 63.5 ± 10.5 | 0.6 ± 0.4 | 15.8 ± 14.2 | -15.2 ± 14.1 |
| | | 298 | 7.3 ± 0.2 | -56.6 ± 7.5 | 64.0 ± 7.5 | 0.1 ± 0.4 | -9.2 ± 12.6 | 9.3 ± 12.7 |
| | | 303 | 8.2 ± 0.4 | -55.1 ± 13.5 | 63.4 ± 13.7 | 0.7 ± 0.5 | -34.9 ± 24.0 | 35.6 ± 24.5 |
| **R188A** | **K243A** | 288 | 1.0 ± 0.2 | 15.7 ± 13.1 | -14.6 ± 12.9 | 3.2 ± 0.5 | -15.1 ± 23.8 | 18.3 ± 23.4 |
| | | 293 | 0.9 ± 0.1 | 0.3 ± 6.2 | 0.6 ± 6.2 | 3.4 ± 0.2 | 2.2 ± 11.1 | 1.2 ± 11.1 |
| | | 298 | 1.1 ± 0.2 | -14.9 ± 6.0 | 16.0 ± 6.2 | 3.2 ± 0.4 | 19.1 ± 11.5 | -16.0 ± 11.8 |
| | | 303 | 1.4 ± 0.4 | -29.9 ± 12.7 | 31.4 ± 13.0 | 2.8 ± 0.6 | 36.0 ± 24.2 | -33.2 ± 24.7 |
| **R188A** | **K299A** | 288 | 7.2 ± 0.2 | 20.6 ± 20.0 | -13.4 ± 19.7 | -1.5 ± 0.5 | 6.2 ± 28.8 | -7.7 ± 28.4 |
| | | 293 | 6.9 ± 0.1 | 25.2 ± 7.5 | -18.3 ± 7.5 | -1.4 ± 0.2 | -14.4 ± 11.9 | 13.0 ± 11.9 |
| | | 298 | 6.6 ± 0.2 | 30.2 ± 9.6 | -23.6 ± 9.6 | -1.1 ± 0.4 | -35.5 ± 13.2 | 34.3 ± 13.3 |
| | | 303 | 6.1 ± 0.4 | 35.3 ± 23.3 | -29.2 ± 23.5 | -0.3 ± 0.6 | -56.8 ± 31.0 | 56.5 ± 31.5 |
| **R238A** | **K243A** | 288 | 3.4 ± 0.2 | -6.4 ± 14.3 | 9.8 ± 14.1 | 1.1 ± 0.5 | -36.9 ± 23.3 | 38.0 ± 22.8 |
| | | 293 | 3.6 ± 0.1 | -51.1 ± 8.1 | 54.7 ± 8.1 | 1.8 ± 0.2 | 1.5 ± 12.4 | 0.3 ± 12.4 |
| | | 298 | 5.6 ± 0.2 | -93.4 ± 7.0 | 99.0 ± 7.2 | 0.8 ± 0.4 | 37.7 ± 12.4 | -37.0 ± 12.7 |
| | | 303 | 7.3 ± 0.4 | -134.4 ± 12.5 | 141.6 ± 12.9 | 0.2 ± 0.6 | 72.8 ± 23.4 | -72.6 ± 24.0 |

**Supplementary Table 4. Triple mutant cycle analysis of the first KDL binding to wild-type and mutant MsbA**. Shown as described in Supplementary Table 3.

| | | | Temperature (K) | ΔΔG (kJ/mol) | ΔΔH (kJ/mol) | Δ(-TΔS) (kJ/mol) | $ΔΔG_{int}$ (kJ/mol) | $ΔΔH_{int}$ (kJ/mol) | $Δ(-ΔTS)_{int}$ (kJ/mol) |
|---|---|---|---|---|---|---|---|---|---|
| R188A | R238A | | 288 | 2.2 ± 0.2 | -70.4 ± 8.9 | 72.6 ± 8.7 | | | |
| | | | 293 | 3.2 ± 0.1 | -62.7 ± 5.4 | 65.9 ± 5.3 | | | |
| | | | 298 | 4.7 ± 0.1 | -53.7 ± 6.1 | 58.3 ± 6.1 | | | |
| | | | 303 | 5.4 ± 0.1 | -43.9 ± 10.2 | 49.3 ± 10.3 | | | |
| | K243A | | 288 | -1.8 ± 0.2 | 2.6 ± 11.0 | 19.9 ± 10.8 | | | |
| | | | 293 | -1.8 ± 0.2 | -4.9 ± 5.0 | -6.4 ± 5.0 | | | |
| | | | 298 | -1.6 ± 0.1 | -11.9 ± 5.4 | 10.4 ± 5.4 | | | |
| | | | 303 | -1.4 ± 0.2 | -18.7 ± 11.5 | 17.3 ± 11.8 | | | |
| | R238A | K243A | 288 | 5.3 ± 0.4 | -2.2 ± 10.0 | 7.5 ± 9.7 | -4.9 ± 0.5 | -65.6 ± 17.4 | 85.0 ± 16.9 |
| | | | 293 | 5.4 ± 0.4 | -2.9 ± 5.1 | 8.4 ± 4.9 | -4.0 ± 0.5 | -64.7 ± 8.9 | 51.2 ± 8.8 |
| | | | 298 | 5.7 ± 0.2 | -3.2 ± 3.8 | 8.9 ± 3.7 | -2.6 ± 0.2 | -62.4 ± 8.9 | 59.8 ± 8.9 |
| | | | 303 | 5.8 ± 0.2 | -3.2 ± 8.9 | 9.0 ± 8.9 | -1.8 ± 0.4 | -59.5 ± 17.8 | 57.7 ± 18.0 |
| R238A | R188A | | 288 | 1.9 ± 0.2 | -26.5 ± 8.9 | 28.4 ± 8.7 | | | |
| | | | 293 | 2.1 ± 0.1 | -10.6 ± 5.4 | 12.8 ± 5.3 | | | |
| | | | 298 | 2.5 ± 0.1 | 6.3 ± 6.1 | -3.7 ± 6.1 | | | |
| | | | 303 | 2.2 ± 0.1 | 23.7 ± 10.2 | -21.5 ± 10.3 | | | |
| | K243A | | 288 | 0.3 ± 0.2 | 24.4 ± 8.0 | -24.1 ± 7.8 | | | |
| | | | 293 | -0.2 ± 0.1 | -4.2 ± 5.3 | 4.0 ± 5.3 | | | |
| | | | 298 | 0.8 ± 0.1 | -30.5 ± 6.2 | 31.3 ± 6.2 | | | |
| | | | 303 | 1.2 ± 0.2 | -55.5 ± 9.9 | 56.7 ± 10.2 | | | |
| | R188A | K243A | 288 | 5.1 ± 0.2 | 41.7 ± 6.6 | -36.6 ± 6.4 | -2.9 ± 0.4 | -43.8 ± 13.7 | 40.9 ± 13.3 |
| | | | 293 | 4.3 ± 0.4 | 49.1 ± 5.4 | -44.8 ± 5.1 | -2.4 ± 0.4 | -64.0 ± 9.3 | 61.6 ± 9.1 |
| | | | 298 | 3.6 ± 0.2 | 56.7 ± 4.9 | -53.2 ± 4.9 | -0.2 ± 0.2 | -81.0 ± 10.0 | 80.8 ± 10.0 |
| | | | 303 | 2.6 ± 0.2 | 64.5 ± 6.7 | -61.9 ± 6.9 | 0.8 ± 0.4 | -96.3 ± 15.7 | 97.1 ± 16.0 |
| K243A | R188A | | 288 | -0.4 ± 0.2 | 28.1 ± 11.0 | -28.5 ± 10.8 | | | |
| | | | 293 | -0.7 ± 0.2 | 3.0 ± 5.0 | -3.7 ± 5.0 | | | |
| | | | 298 | -0.5 ± 0.1 | -22.1 ± 5.4 | 21.6 ± 5.4 | | | |
| | | | 303 | 0.1 ± 0.2 | -47.2 ± 11.5 | 47.3 ± 11.8 | | | |
| | R238A | | 288 | 1.9 ± 0.2 | 6.1 ± 8.0 | -4.1 ± 7.8 | | | |
| | | | 293 | 2.0 ± 0.1 | -48.4 ± 5.3 | 50.4 ± 5.3 | | | |
| | | | 298 | 4.0 ± 0.1 | -100.6 ± 6.2 | 104.6 ± 6.2 | | | |
| | | | 303 | 5.9 ± 0.2 | -151.6 ± 9.9 | 157.5 ± 10.2 | | | |
| | R188A | R238A | 288 | 6.7 ± 0.2 | 23.3 ± 9.2 | -16.6 ± 8.9 | -5.2 ± 0.4 | 10.9 ± 16.4 | -16.0 ± 16.0 |
| | | | 293 | 6.6 ± 0.4 | 5.0 ± 5.1 | 1.6 ± 4.9 | -5.2 ± 0.5 | -50.4 ± 8.9 | 45.1 ± 8.8 |
| | | | 298 | 6.8 ± 0.2 | -13.3 ± 3.9 | 20.1 ± 4.0 | -3.3 ± 0.2 | -109.4 ± 9.2 | 106.1 ± 9.2 |
| | | | 303 | 7.3 ± 0.2 | -31.6 ± 8.7 | 38.9 ± 8.9 | -1.3 ± 0.4 | -167.2 ± 17.5 | 165.9 ± 17.9 |

**Supplementary table 5. Equilibrium dissociation constants ($K_D$) for KDL binding MsbA trapped with ADP and vanadate at various temperatures.** Reported are the mean and standard deviation (*n* = 3)

| | Temperature(K) | $K_{D1}$(μM) | $K_{D2}$(μM) | $K_{D3}$(μM) | $R^{2*}$ | $X^{2*}$ |
|---|---|---|---|---|---|---|
| **WT** | 293 | 0.51 ± 0.04 | 1.16 ± 0.07 | | 0.97 | 0.08 |
| | 298 | 0.44 ± 0.02 | 0.93 ± 0.09 | | 0.94 | 0.17 |
| | 303 | 0.38 ± 0.02 | 0.73 ± 0.05 | | 0.94 | 0.15 |
| | 310 | 0.31 ± 0.01 | 0.53 ± 0.04 | | 0.92 | 0.2 |
| **R188A** | 293 | 1.56 ± 0.09 | 3.46 ± 0.15 | 10.98 ± 1.15 | 0.98 | 0.07 |
| | 298 | 1.53 ± 0.11 | 3.40 ± 0.21 | 10.15 ± 1.62 | 0.98 | 0.06 |
| | 303 | 1.35 ± 0.10 | 2.90 ± 0.26 | 9.02 ± 1.11 | 0.98 | 0.05 |
| | 310 | 0.93 ± 0.09 | 1.89 ± 0.24 | 5.50 ± 1.10 | 0.97 | 0.07 |
| **R238A** | 293 | 1.67 ± 0.23 | 6.71 ± 1.38 | | 0.99 | 0.05 |
| | 298 | 0.91 ± 0.10 | 3.15 ± 0.12 | | 0.98 | 0.06 |
| | 303 | 0.34 ± 0.04 | 1.09 ± 0.23 | 3.75 ± 0.60 | 0.95 | 0.11 |
| | 310 | 0.10 ± 0.02 | 0.33 ± 0.11 | 0.92 ± 0.38 | 1 | 0 |
| **K243A** | 293 | 2.01 ± 0.06 | 5.26 ± 0.85 | | 0.99 | 0.02 |
| | 298 | 1.46 ± 0.01 | 3.90 ± 0.48 | | 0.99 | 0.03 |
| | 303 | 0.60 ± 0.02 | 1.54 ± 0.12 | 3.72 ± 0.54 | 0.98 | 0.04 |
| | 310 | 0.25 ± 0.02 | 0.46 ± 0.02 | 1.06 ± 0.04 | 0.89 | 0.2 |
| **R188A R238A** | 293 | 1.24 ± 0.10 | 4.82 ± 0.16 | | 0.99 | 0.03 |
| | 298 | 1.17 ± 0.11 | 4.65 ± 0.92 | | 0.99 | 0.02 |
| | 303 | 0.87 ± 0.06 | 3.05 ± 0.26 | | 0.98 | 0.04 |
| | 310 | 0.39 ± 0.05 | 1.28 ± 0.09 | 6.36 ± 1.07 | 0.98 | 0.04 |
| **R188A K243A** | 293 | 3.64 ± 0.28 | 15.66 ± 4.76 | | 0.99 | 0.03 |
| | 298 | 2.23 ± 0.07 | 6.68 ± 0.17 | | 0.99 | 0.03 |
| | 303 | 1.06 ± 0.05 | 3.10 ± 0.33 | 19.33 ± 3.70 | 0.99 | 0.03 |
| | 310 | 0.66 ± 0.02 | 1.56 ± 0.04 | 5.70 ± 0.93 | 0.98 | 0.04 |
| **R238A K243A** | 293 | 1.31 ± 0.04 | 4.35 ± 0.07 | | 0.99 | 0.03 |
| | 298 | 1.03 ± 0.09 | 3.65 ± 0.15 | | 0.99 | 0.02 |
| | 303 | 0.48 ± 0.05 | 1.97 ± 0.09 | | 0.97 | 0.09 |
| | 310 | 0.10 ± 0.02 | 0.58 ± 0.06 | | 0.74 | 0.93 |

*These values represent the replicates with the poorest fits.

**Supplementary table 6. Thermodynamic signatures of KDL interacting with wild-type and mutant MsbA trapped with ADP and vanadate.** Reported are the mean with standard deviation ($n$ = 3), and the subscript denotes the $n^{th}$ KDL binding event.

| | T (K) | $\Delta G_1$ (kJ/mol) | $\Delta H_1$ (kJ/mol) | $-T\Delta S_1$ (kJ/mol) | $\Delta Cp_1$ (kJ/mol/K) | $\Delta G_2$ (kJ/mol) | $\Delta H_2$ (kJ/mol) | $-T\Delta S_2$ (kJ/mol) | $\Delta Cp_2$ (kJ/mol/K) | $\Delta G_3$ (kJ/mol) | $\Delta H_3$ (kJ/mol) | $-T\Delta S_3$ (kJ/mol) | $\Delta Cp_3$ (kJ/mol/K) |
|---|---|---|---|---|---|---|---|---|---|---|---|---|---|
| **WT** | 293 | -35.3 ± 0.1 | 21.9 ± 1.5 | -57.2 ± 1.3 | | -33.3 ± 0.2 | 35.0 ± 0.6 | -68.3 ± 0.6 | | | | | |
| | 298 | -36.3 ± 0.1 | 21.9 ± 1.5 | -58.2 ± 1.3 | | -34.4 ± 0.2 | 35.0 ± 0.6 | -69.4 ± 0.6 | | | | | |
| | 303 | -37.3 ± 0.1 | 21.9 ± 1.5 | -59.2 ± 1.3 | | -35.6 ± 0.2 | 35.0 ± 0.6 | -70.6 ± 0.6 | | | | | |
| | 310 | -38.6 ± 0.1 | 21.9 ± 1.5 | -60.5 ± 1.3 | | -37.2 ± 0.2 | 35.0 ± 0.6 | -72.2 ± 0.6 | | | | | |
| **R188A** | 293 | -32.6 ± 0.1 | -6.6 ± 2.6 | -26.0 ± 2.7 | 3.6 ± 0.1 | -30.7 ± 0.1 | -7.2 ± 2.4 | -23.4 ± 2.5 | 4.1 ± 0.2 | -27.8 ± 0.3 | -3.2 ± 21.3 | -24.7 ± 21.1 | 4.1 ± 2.7 |
| | 298 | -33.2 ± 0.2 | 11.3 ± 2.6 | -44.5 ± 2.7 | 3.5 ± 0.1 | -31.2 ± 0.1 | 13.4 ± 3.3 | -44.7 ± 3.4 | 4.2 ± 0.3 | -28.5 ± 0.4 | 17.6 ± 12.4 | -46.1 ± 12.3 | 3.6 ± 4.2 |
| | 303 | -34.1 ± 0.2 | 29.2 ± 2.5 | -63.3 ± 2.6 | 3.6 ± 0.1 | -32.2 ± 0.2 | 34.1 ± 4.3 | -66.3 ± 4.5 | 4.1 ± 0.2 | -29.3 ± 0.3 | 37.8 ± 14.8 | -67.1 ± 15.1 | 4.9 ± 1.4 |
| | 310 | -35.8 ± 0.2 | 54.3 ± 2.6 | -90.2 ± 2.8 | 3.6 ± 0.1 | -34.0 ± 0.3 | 63.0 ± 5.8 | -97.0 ± 6.1 | 4.1 ± 0.2 | -31.3 ± 0.5 | 67.9 ± 26.1 | -99.2 ± 26.6 | 4.3 ± 2.2 |
| **R238A** | 293 | -32.1 ± 0.2 | 126.8 ± 5.8 | -158.9 ± 5.9 | | -28.9 ± 0.4 | 137.7 ± 10.8 | -166.5 ± 10.8 | | | | | |
| | 298 | -34.8 ± 0.4 | 126.8 ± 5.8 | -161.6 ± 6.0 | | -31.7 ± 0.4 | 137.7 ± 10.8 | -169.4 ± 11.0 | | | | | |
| | 303 | -37.6 ± 0.4 | 126.8 ± 5.8 | -164.3 ± 6.1 | | -34.6 ± 0.5 | 137.7 ± 10.8 | -172.2 ± 11.1 | | | | | |
| | 310 | -41.4 ± 0.5 | 126.8 ± 5.8 | -168.1 ± 6.2 | | -38.5 ± 0.7 | 137.7 ± 10.8 | -176.2 ± 11.4 | | | | | |
| **K243A** | 293 | -31.6 ± 0.1 | 96.2 ± 5.1 | -127.9 ± 5.1 | | -29.1 ± 0.2 | 111.7 ± 6.9 | -140.8 ± 6.6 | | | | | |
| | 298 | -33.8 ± 0.1 | 96.2 ± 5.1 | -130.0 ± 5.1 | | -31.5 ± 0.1 | 111.7 ± 6.9 | -143.2 ± 6.7 | | | | | |
| | 303 | -36.0 ± 0.1 | 96.2 ± 5.1 | -132.2 ± 5.3 | | -33.9 ± 0.1 | 111.7 ± 6.9 | -145.6 ± 6.9 | | | | | |
| | 310 | -39.1 ± 0.2 | 96.2 ± 5.1 | -135.3 ± 5.4 | | -37.3 ± 0.2 | 111.7 ± 6.9 | -149 ± 7.1 | | | | | |
| **R188A R238A** | 293 | -33.2 ± 0.2 | -10.2 ± 3.0 | -23.0 ± 3.1 | 7.5 ± 1.0 | -29.8 ± 0.1 | -8.2 ± 30.4 | -21.7 ± 30.3 | 8.1 ± 3.5 | | | | |
| | 298 | -33.9 ± 0.2 | 27.2 ± 2.5 | -61.0 ± 2.6 | 7.5 ± 0.3 | -30.5 ± 0.5 | 31.9 ± 13.7 | -62.4 ± 14.2 | 8.6 ± 4.6 | | | | |
| | 303 | -35.2 ± 0.2 | 64.5 ± 7.3 | -99.7 ± 7.2 | 7.5 ± 2.1 | -32.0 ± 0.2 | 72.5 ± 4.9 | -104.5 ± 4.8 | 7.4 ± 1.9 | | | | |
| | 310 | -38.1 ± 0.4 | 116.9 ± 16.4 | -155.0 ± 16.8 | 7.5 ± 1.3 | -35.0 ± 0.2 | 127.7 ± 25.9 | -162.7 ± 25.9 | 7.9 ± 3.0 | | | | |
| **R188A K243A** | 293 | -30.5 ± 0.2 | 92.4 ± 8.6 | -122.9 ± 8.4 | -1.8 ± 0.9 | -27.0 ± 0.7 | 135.4 ± 44.8 | -162.4 ± 44.1 | -4.0 ± 4.0 | | | | |
| | 298 | -32.3 ± 0.1 | 82.1 ± 4.4 | -114.4 ± 4.3 | -0.1 ± 0.9 | -29.5 ± 0.1 | 114.7 ± 25.2 | -144.2 ± 25.2 | -3.4 ± 4.5 | | | | |
| | 303 | -34.7 ± 0.1 | 73.4 ± 0.1 | -108.0 ± 0.1 | -4.4 ± 0.8 | -32.0 ± 0.3 | 94.5 ± 5.7 | -126.5 ± 5.6 | -4.9 ± 3.8 | | | | |
| | 310 | -36.7 ± 0.1 | 55.6 ± 5.7 | -92.3 ± 5.7 | -2.6 ± 0.8 | -34.5 ± 0.1 | 64.4 ± 22.9 | -98.9 ± 22.9 | -4.3 ± 3.9 | | | | |
| **R238A K243A** | 293 | -33.0 ± 0.1 | 8.9 ± 14.2 | -41.9 ± 14.2 | 12.7 ± 2.7 | -30.1 ± 0.1 | 6.5 ± 2.2 | -36.6 ± 2.1 | 10.0 ± 0.7 | | | | |
| | 298 | -34.2 ± 0.2 | 71.8 ± 7.2 | -106.0 ± 7.4 | 13.2 ± 2.1 | -31.0 ± 0.1 | 56.1 ± 2.1 | -87.1 ± 2.2 | 10.5 ± 0.9 | | | | |
| | 303 | -36.7 ± 0.2 | 135.3 ± 16.2 | -171.9 ± 16.3 | 11.9 ± 3.5 | -33.1 ± 0.1 | 106.2 ± 5.4 | -139.3 ± 5.5 | 9.1 ± 1.0 | | | | |
| | 310 | -41.6 ± 0.8 | 222.4 ± 35.8 | -264.0 ± 36.5 | 12.4 ± 2.9 | -37.0 ± 0.3 | 174.4 ± 10.5 | -211.4 ± 10.8 | 9.7 ± 0.7 | | | | |

**Supplementary Table 7. Double mutant cycle analysis of the first KDL binding to wild-type and mutant MsbA trapped with ADP and vanadate**. Shown as described in Supplementary Table 3.

| | | Temperature (K) | ΔΔG (kJ/mol) | ΔΔH (kJ/mol) | Δ(-TΔS) (kJ/mol) | $ΔΔG_{int}$ (kJ/mol) | $ΔΔH_{int}$ (kJ/mol) | $Δ(-ΔTS)_{int}$ (kJ/mol) |
|---|---|---|---|---|---|---|---|---|
| **R188A** | | 293 | 2.7 ± 0.1 | -28.5 ± 3.1 | 31.2 ± 3.1 | | | |
| | | 298 | 3.1 ± 0.2 | -10.6 ± 2.9 | 13.7 ± 3.1 | | | |
| | | 303 | 3.2 ± 0.2 | 7.3 ± 2.9 | -4.1 ± 2.9 | | | |
| | | 310 | 2.8 ± 0.2 | 32.4 ± 2.9 | -29.7 ± 3.1 | | | |
| **R238A** | | 293 | 3.2 ± 0.2 | 104.9 ± 6.0 | -101.7 ± 6.0 | | | |
| | | 298 | 1.5 ± 0.4 | 104.9 ± 6.0 | -103.4 ± 6.1 | | | |
| | | 303 | -0.3 ± 0.4 | 104.9 ± 6.0 | -105.1 ± 6.2 | | | |
| | | 310 | -2.8 ± 0.5 | 104.9 ± 6.0 | -107.6 ± 6.4 | | | |
| **K243A** | | 293 | 3.7 ± 0.1 | 74.3 ± 5.4 | -70.7 ± 5.3 | | | |
| | | 298 | 2.5 ± 0.1 | 74.3 ± 5.4 | -71.8 ± 5.3 | | | |
| | | 303 | 1.3 ± 0.1 | 74.3 ± 5.4 | -73.0 ± 5.4 | | | |
| | | 310 | -0.5 ± 0.2 | 74.3 ± 5.4 | -74.8 ± 5.5 | | | |
| **R188A** | **R238A** | 293 | 2.2 ± 0.2 | -32.1 ± 3.3 | 34.2 ± 3.4 | 3.8 ± 0.4 | 108.5 ± 7.0 | -104.7 ± 7.2 |
| | | 298 | 2.4 ± 0.2 | 5.3 ± 2.9 | -2.8 ± 2.9 | 2.2 ± 0.5 | 89.0 ± 6.9 | -86.9 ± 7.1 |
| | | 303 | 2.1 ± 0.2 | 42.6 ± 7.5 | -40.5 ± 7.3 | 0.8 ± 0.5 | 69.6 ± 9.6 | -68.7 ± 9.8 |
| | | 310 | 0.5 ± 0.4 | 95.0 ± 16.5 | -94.5 ± 16.8 | -0.5 ± 0.6 | 42.4 ± 17.6 | -42.8 ± 18.1 |
| **R188A** | **K243A** | 293 | 4.8 ± 0.2 | 70.5 ± 8.7 | -65.7 ± 8.6 | 1.6 ± 0.2 | -24.7 ± 10.4 | 26.3 ± 10.3 |
| | | 298 | 4.0 ± 0.1 | 60.2 ± 4.7 | -56.2 ± 4.5 | 1.6 ± 0.2 | 3.5 ± 7.2 | -1.9 ± 7.2 |
| | | 303 | 2.6 ± 0.1 | 51.5 ± 1.5 | -48.8 ± 1.3 | 1.9 ± 0.2 | 30.1 ± 5.8 | -28.2 ± 5.9 |
| | | 310 | 1.9 ± 0.1 | 33.7 ± 5.9 | -31.8 ± 5.9 | 0.4 ± 0.4 | 73.0 ± 8.1 | -72.7 ± 8.3 |
| **R238A** | **K243A** | 293 | 2.3 ± 0.1 | -13.0 ± 14.3 | 15.3 ± 14.2 | 4.6 ± 0.2 | 192.2 ± 16.2 | -187.7 ± 16.2 |
| | | 298 | 2.1 ± 0.2 | 49.9 ± 7.3 | -47.8 ± 7.6 | 1.9 ± 0.4 | 129.3 ± 10.5 | -127.4 ± 10.9 |
| | | 303 | 0.6 ± 0.2 | 113.4 ± 16.3 | -112.7 ± 16.4 | 0.4 ± 0.5 | 65.8 ± 17.9 | -65.4 ± 18.2 |
| | | 310 | -3.0 ± 0.9 | 200.5 ± 35.8 | -203.5 ± 36.5 | -0.4 ± 1.0 | -21.3 ± 36.6 | 21.1 ± 37.5 |

**Supplementary Table 8**. **Double mutant cycle analysis of the second KDL binding to wild-type and mutant MsbA trapped with ADP and vanadate**. Shown as described in Supplementary Table 3.

| | | Temperature (K) | ΔΔG (kJ/mol) | ΔΔH (kJ/mol) | Δ(-TΔS) (kJ/mol) | $\Delta\Delta G_{int}$ (kJ/mol) | $\Delta\Delta H_{int}$ (kJ/mol) | $\Delta(-\Delta TS)_{int}$ (kJ/mol) |
|---|---|---|---|---|---|---|---|---|
| R188A | | 293 | 2.7 ± 0.2 | -42.2 ± 2.4 | 44.9 ± 2.6 | | | |
| | | 298 | 3.2 ± 0.2 | -21.6 ± 3.3 | 24.8 ± 3.4 | | | |
| | | 303 | 3.5 ± 0.4 | -0.9 ± 4.4 | 4.3 ± 4.5 | | | |
| | | 310 | 3.2 ± 0.4 | 28.0 ± 5.9 | -24.8 ± 6.1 | | | |
| R238A | | 293 | 4.4 ± 0.5 | 102.7 ± 10.8 | -98.2 ± 10.8 | | | |
| | | 298 | 2.7 ± 0.5 | 102.7 ± 10.8 | -100.0 ± 11.0 | | | |
| | | 303 | 1.0 ± 0.5 | 102.7 ± 10.8 | -101.6 ± 11.1 | | | |
| | | 310 | -1.3 ± 0.7 | 102.7 ± 10.8 | -104.0 ± 11.4 | | | |
| K243A | | 293 | 4.2 ± 0.4 | 76.7 ± 6.9 | -72.5 ± 6.6 | | | |
| | | 298 | 2.9 ± 0.2 | 76.7 ± 6.9 | -73.8 ± 6.7 | | | |
| | | 303 | 1.7 ± 0.2 | 76.7 ± 6.9 | -75.0 ± 6.9 | | | |
| | | 310 | -0.1 ± 0.4 | 76.7 ± 6.9 | -76.8 ± 7.1 | | | |
| R188A | R238A | 293 | 3.5 ± 0.2 | -43.2 ± 30.4 | 46.7 ± 30.3 | 3.6 ± 0.4 | 103.7 ± 32.3 | -100.0 ± 32.2 |
| | | 298 | 3.9 ± 0.5 | -3.1 ± 13.7 | 7.0 ± 14.2 | 2.0 ± 0.6 | 84.2 ± 17.8 | -82.3 ± 18.2 |
| | | 303 | 3.6 ± 0.4 | 37.5 ± 4.9 | -33.9 ± 4.8 | 0.9 ± 0.6 | 64.3 ± 12.6 | -63.3 ± 13.0 |
| | | 310 | 2.2 ± 0.2 | 92.7 ± 25.8 | -90.5 ± 25.8 | -0.3 ± 0.9 | 38.0 ± 28.7 | -38.3 ± 28.9 |
| R188A | K243A | 293 | 6.3 ± 0.7 | 100.4 ± 44.8 | -94.1 ± 44.1 | 0.6 ± 0.7 | -65.9 ± 45.4 | 66.5 ± 44.7 |
| | | 298 | 4.9 ± 0.2 | 79.7 ± 25.2 | -74.8 ± 25.2 | 1.2 ± 0.2 | -24.6 ± 26.3 | 25.8 ± 26.3 |
| | | 303 | 3.6 ± 0.4 | 59.5 ± 5.8 | -55.9 ± 5.6 | 1.5 ± 0.4 | 16.3 ± 9.9 | -14.7 ± 9.9 |
| | | 310 | 2.7 ± 0.2 | 29.4 ± 22.9 | -26.7 ± 22.9 | 0.4 ± 0.4 | 75.3 ± 24.6 | -74.9 ± 24.7 |
| R238A | K243A | 293 | 3.2 ± 0.2 | -28.5 ± 2.2 | 31.7 ± 2.2 | 5.4 ± 0.5 | 207.9 ± 13.0 | -202.4 ± 12.9 |
| | | 298 | 3.4 ± 0.2 | 21.1 ± 2.2 | -17.7 ± 2.3 | 2.2 ± 0.4 | 158.3 ± 13.0 | -156.1 ± 13.1 |
| | | 303 | 2.5 ± 0.2 | 71.2 ± 5.4 | -68.7 ± 5.5 | 0.2 ± 0.5 | 108.2 ± 13.8 | -107.9 ± 14.2 |
| | | 310 | 0.2 ± 0.4 | 139.4 ± 10.5 | -139.2 ± 10.8 | -1.6 ± 0.9 | 40.0 ± 16.5 | -41.6 ± 17.3 |
